# Perinuclear anchoring of telomeres enables plant infection by *Ustilago maydis*

**DOI:** 10.64898/2026.04.03.716281

**Authors:** Estela Sanz-Marti, Manuel Velasco-Gomariz, Blanca Navarrete, José Ignacio Ibeas, Ramón R. Barrales

## Abstract

The spatial organization of the genome is critical for cellular function, yet its contribution to the success of host-pathogen interactions remains poorly understood. A conserved hallmark of nuclear architecture is the perinuclear localization of heterochromatin, which includes not only specific chromosomal subdomains but also major structural elements like centromeres and telomeres. To determine the importance of this organization during fungal pathogenesis, we used the maize pathogen *Ustilago maydis* as an infection model system. We identify the lamina-associated protein homolog, Lem2, as a molecular tether for telomeres and show its deletion abrogates fungal penetration, causing developmental arrest at the appressorium stage. Mechanistically, this infection defect is associated with a failure in nuclear migration. Using hyphal filaments induced under axenic conditions, we observed that mutant nuclei consistently fail to transit from the mother cell into the growing filament. Furthermore, RNA-seq analysis correlates this failure with a marked misregulation of the DNA damage response (DDR) and cell cycle control, including the aberrant expression of multiple checkpoint and signaling proteins. Crucially, artificially tethering telomeres to the nuclear periphery in this mutant partially restores the nuclear migration defect and plant penetration, while simultaneously suppressing the aberrant DDR. This work establishes a functional link between a specific spatial nuclear configuration and fungal infectivity, revealing the conserved telomere-anchoring machinery as a potential target for novel antifungal strategies.

## Introduction

In eukaryotes, DNA is packaged around histones to form chromatin. This packaging is not uniform; post-translational modifications of histones and DNA modifications create different levels of accessibility, primarily distinguishing transcriptionally active euchromatin from transcriptionally silent heterochromatin. A key feature of nuclear organization is the tethering of heterochromatic domains, known as Lamina-Associated Domains (LADs), to the nuclear lamina (1). This peripheral microenvironment is thought to facilitate the establishment, propagation, and maintenance of a repressed transcriptional state (2). The binding of silenced chromatin to the nuclear periphery is dynamic, playing a crucial role in processes like cell differentiation and development (3–6). Beyond the positioning of these specific domains, the tethering of silent chromosomal regions to the nuclear periphery is an evolutionarily conserved phenomenon. For instance, telomeres adopt a perinuclear position in organisms from yeast to mammals (7–12) and centromeres show a comparable localization in yeast, flies, and plants (8, 13).

The nuclear lamina is primarily responsible for such chromatin anchoring through its components: the A- and B-type lamins and the integral inner nuclear membrane proteins (14). The latter group includes several key lamina-associated proteins (LAPs) that tether silent chromatin, most notably the Lamin B receptor (LBR) and the LAP2, emerin, MAN1 (LEM)-domain family of proteins (15, 16). However, dissecting the specific contribution of these proteins in metazoans is challenging due to a complex network of proteins with interdependent localization (17–19). Thus, many studies have utilized yeast, primarily *Saccharomyces cerevisiae* and *Schizosaccharomyces pombe*, to investigate the molecular basis of chromatin tethering. While these lower eukaryotes lack a nuclear lamina (20), their genomes encode functional LAP analogs, such as the LEM-like domain containing proteins Lem2 and Man1 in *S. pombe* (21, 22). These proteins are critical for nuclear envelope structure and have conserved roles in anchoring heterochromatin, including centromeres and telomeres, to the nuclear periphery to ensure transcriptional silencing (22–30).

The functional significance of this peripheral anchoring extends beyond static gene silencing and involves highly dynamic processes. On one hand, LADs exhibit multi-level dynamic organization, including stimulus-driven shifts and developmental restructuring to control cell-specific genes (6). On the other hand, the nuclear periphery also serves as a critical signaling hub, particularly for pathways that monitor genome integrity, such as the DNA Damage Response (DDR). In human cells, for instance, the choice of repair system for a double-strand break depends on its nuclear location. Breaks induced at the nuclear membrane are repaired by alternative end-joining, while those occurring in nuclear pores or the nuclear interior are repaired using homologous recombination (31). In addition, specific lamins and LAPs are involved in the recruitment of key DNA damage and checkpoint proteins to the nuclear envelope in metazoans (32–35). Tethering of telomeres to the nuclear envelope, in particular, affects double-strand break repair by reducing homology search efficiency (36, 37). In addition to their implication in DDR, different LEM domain containing proteins have been described to be involved in the regulation of various signaling pathways such as the Wnt, Notch and Rb/MyoD pathways (38–40), TGF-ß and bone morphogenic protein (BMP) (41–43), and regulation of MAP and AKT kinases (44).

While LAPs are well established as key signaling hubs for genome integrity and cell development, their contribution to the drastic cellular reprogramming and morphogenetic events that must be precisely monitored and controlled during fungal pathogenesis represents a significant gap in our knowledge. The maize pathogen *Ustilago maydis* is a paradigm model system to study these processes (45). Its dimorphic behavior, the subsequent morphological changes produced during infection, and the genetic and cellular biology tools available make it an exceptional system in which to investigate the role of LAPs during infection.

*U. maydis* belongs to the smut fungi that widely affect crop cultivars (46). The infection cycle begins when two sexually compatible yeast-like cells recognize each other and initiate filamentation (47, 48). These conjugation tubes then fuse, forming a dikaryotic filament in which the two nuclei remain unfused and arrested in the G2 phase of the cell cycle (49, 50). This dikaryotic filament proliferates on the plant surface, culminating in the formation of a specialized penetration structure, the appressorium (51–53). Once this swollen hyphal tip invades the plant, the cell cycle block is released and a proliferative, septated hypha colonizes the host tissue (54). Finally, the fungus induces tumors on the plant where spores are produced. Following the release and germination of these spores, the cycle starts again (55). This pathogenic program is tightly controlled by a complex regulatory network strictly coupled to sexual development. Following pheromone recognition, the activation of the cAMP/PKA and MAP kinase pathways leads to the expression of the compatible mating-type genes bE and bW. Together, they form a heterodimer that acts as the master transcriptional regulator, orchestrating a downstream hierarchical cascade essential for successful host colonization (56).

To investigate the role of LAPs during the morphological and regulatory changes of fungal pathogenesis, we identified and deleted the sole LEM-domain protein in the *U. maydis* genome, hereafter Lem2. We demonstrate that Lem2 is essential for plant infection by functioning as the molecular tether for telomeres. We show that this anchoring provides a crucial regulatory signal required for nuclear migration during the formation of the infective hypha, thereby establishing a direct link between telomere positioning and fungal pathogenesis.

## Results

### Lem2 mediates telomere anchoring to the nuclear envelope in *U. maydis*

To investigate the role of lamin-associated proteins in *Ustilago maydis*, we specifically focused on conserved LEM-domain-containing proteins. Sequence analysis identified a single protein containing this domain (UMAG_00208), hereafter referred to as Lem2. This protein is widely conserved across fungi and exhibits around 20% sequence identity with *Schizosaccharomyces pombe* Lem2 and *Saccharomyces cerevisiae* Heh1. As expected, it features the two characteristic Lem2 domains: the MSC and the LEM domains (Fig. 1A). These domains have been well-characterized in *S. pombe*, where the MSC domain plays a critical role in anchoring telomeres to the nuclear envelope (NE) and in heterochromatin silencing, while the LEM domain mediates centromere anchoring (23). We first sought to confirm that *U. maydis* Lem2 localizes to the nuclear periphery, as previously described in other fungal models (22, 23, 57). Live-cell fluorescence microscopy of cells expressing Lem2-GFP at the endogenous locus confirmed that the protein is exclusively localized to the NE (Fig. 1B). Notably, within this peripheral distribution, Lem2-GFP exhibited multiple dynamic foci scattered throughout the NE (Fig. 1B and S1A). This pattern differs from *S. pombe*, where Lem2 concentrates in one or two distinct spots that colocalize with the spindle pole body (SPB) (22, 57). We next investigated whether the role of Lem2 in maintaining nuclear integrity (22, 57) is also conserved in *U. maydis.* To this end, we monitored the leakage of a nuclear reporter (NLS::3xRFP) while simultaneously labeling the NE with the nucleoporin Nup214::GFP.

**Figure 1.**
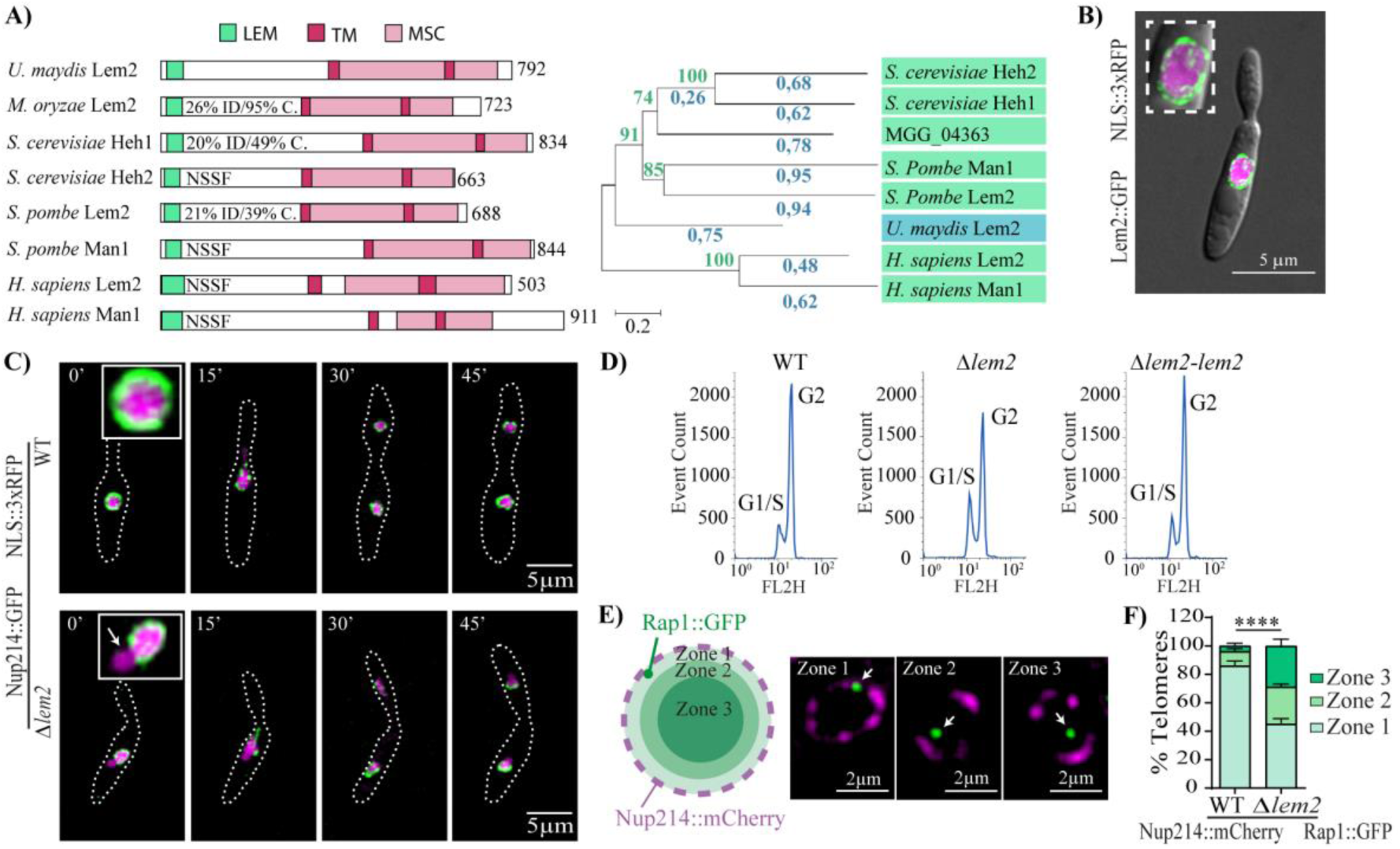
Lem2 is required for nuclear envelope integrity and telomere anchoring. **(A)** Left panel: Schematic representation of Lem2 domain organization in *U. maydis, Magnaporthe oryzae, S. cerevisiae, S. pombe, and Homo sapiens*. Percentage of sequence identity (% ID) and coverage (C.) are indicated for each ortholog; NSSF (no significant similarity found) denotes no significant homology detected. TM, transmembrane domain. Right panel: Phylogenetic analysis of Lem2 orthologs across the same species. Bootstrap values are indicated in green, while evolutionary distances are shown in blue above each branch. **(B)** Live-cell imaging of Lem2::GFP NE localization. NLS::3xRFP (magenta) was used as a nuclear marker. **(C)** Representative time-lapse frames of mitotic wild-type (WT) and *Δlem2* cells. Strains express Nup214::GFP as a NE marker and NLS::3xRFP as a reporter for nuclear integrity. DDotted lines delineate cell boundaries. The white arrow indicates a nuclear protrusion. **(D)** FACS analysis of WT, *Δlem2,* and *Δlem2-lem2* strains. **(E)** Three-zone method. Left: Schematic illustrating the division of the nucleus into three concentric zones (I–III) of equal area. Right: Representative images showing telomeres (Rap1::GFP) localized in Zone I, II, or III. The nuclear envelope is labeled with Nup214::mCherry. Statistical significance was determined using the ordinal logistic mixed model (GLMM) (****, *P* < 0.0001). **(F)** Quantification of Rap1::GFP distribution using the three-zone method. The number of analyzed cells (*n*) is indicated above each column. Error bars represent the standard deviation (SD) from three independent replicates. Images in (C) are maximum intensity projections of Z-stacks. All panels feature strains in the SG200 genetic background.

In the *Δlem2* mutant, the NLS::3xRFP reporter remained nuclear until the physiological breakdown of the nuclear envelope characteristic of its open mitosis, indicating an absence of gross NE rupture. However, the presence of prominent nuclear protrusions in regions lacking Nup214 revealed that nuclear structure is compromised (Fig. 1C). Despite these NE defects, the *Δlem2* mutant completes mitosis normally and shows only a slight reduction in growth rate (Fig. S1B), which may be associated with a moderate accumulation of cells in the G1/S phase (Fig. 1D). Importantly, a recovery of growth capacity and the reversal of cell-cycle defects were observed upon reintroducing *lem2* into the *ip* locus of the *Δlem2* mutant (Fig. S1B and 1D).

Beyond their role in NE integrity, LAPs have emerged as central components responsible for tethering silenced chromatin to the nuclear periphery. Given the conserved role of Lem2 in telomere tethering in other fungi (22), we examined whether *U. maydis* Lem2 plays a similar role in perinuclear telomere positioning. To this end, we analyzed the localization of the telomere-binding protein Rap1 (GFP-tagged) relative to the nucleoporin Nup214::mCherry using the three-zone method (Fig. 1E) (58). Live-cell imaging confirmed that the expression of Rap1::GFP did not affect mitotic progression in either wild-type or *Δlem2* cells (Fig. S1C). In wild-type cells, telomeres were strongly enriched at the nuclear periphery (86% in Zone I, 10% in Zone II, and 4% in Zone III), indicating their anchoring to the NE in *U. maydis*. In contrast, a significant reduction in the peripheral enrichment of telomeres was observed in the *Δlem2* strain (45% in Zone I), demonstrating that Lem2 is required for efficient telomere anchoring (Fig. 1F). Moreover, no changes in centromere positioning or SPB location were observed in the absence of Lem2, as determined by the localization of the inner kinetochore protein Mis12::GFP and the centromere-associated protein Grc1::GFP (Fig. S1D and E). Taken together, these results demonstrate that Lem2 is a core structural component of the *U. maydis* nuclear envelope, playing a specific and conserved role in tethering telomeres to the nuclear periphery.

### Lem2 is essential for fungal penetration of plant tissue

Having established that Lem2 anchors telomeres to the nuclear periphery, we next sought to investigate the potential role of this nuclear organization in virulence. Deletion of *lem2* resulted in a complete loss of virulence in both the solopathogenic (SG200) and sexually compatible (FB1xFB2) backgrounds. Virulence was rescued when *lem2* was reintroduced, confirming that the virulence defect is specifically due to the absence of Lem2 (Fig. 2A).

**Figure 2.**
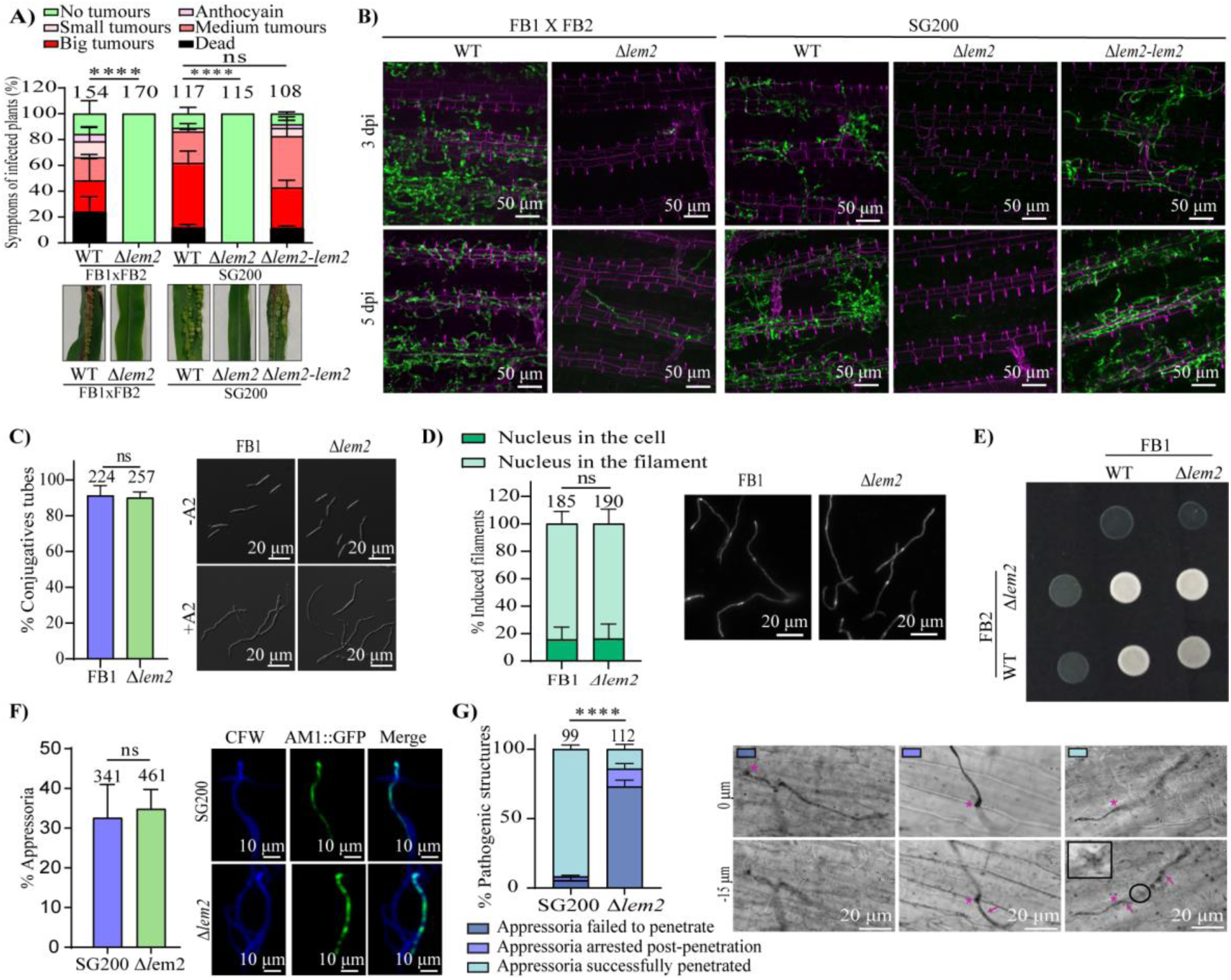
Deletion of *lem2* abrogates plant penetration, leading to a complete loss of virulence in *Ustilago maydis*. **(A)** Disease symptoms in maize plants infected with the indicated strains at 14 days post-inoculation (dpi). The top panel shows the quantification of symptom categories, while the bottom panel displays representative images of the different symptom classes. **(B)** Colonization of maize leaves infected with WT, *Δlem2*, and *Δlem2-*lem2 strains at 3 and 5 dpi. Plant cell walls were visualized with propidium iodide (magenta) and fungal hyphae with WGA-Alexa Fluor 488 (green). Images represent maximum intensity projections of Z-stacks. **(C)** Quantification of conjugation tube formation. Left panel: Percentage of FB1 and FB1 *Δlem2* cells 5 hours post-induction (hpi) with synthetic *a2* pheromone. Right panel: Representative images of FB1 and FB1 *Δlem2* conjugation tubes. **(D)** Quantification of nuclear localization in conjugation tubes. FB1 and FB1 *Δlem2* cells were analyzed at 5 hpi following *a2* pheromone induction. Nuclei were visualized using DAPI staining. Representative images of WT and *Δlem2* conjugation tubes are shown. **(E)** Mating assay of compatible *U. maydis* strains. Cultures of FB1, FB2, FB1 *Δlem2*, and FB2 *Δlem2* were spotted on PD-charcoal plates. **(F)** *In vitro* quantification of appressorium formation. Filaments from the SG200 and SG200 *Δlem2* strains were analyzed for their ability to differentiate into appressoria. The AM1::GFP reporter (green) specifically marks filaments differentiating into appressoria, characterized by a crook-like structure. Fungal cell walls were visualized with calcofluor white (blue). Images represent maximum intensity projections of Z-stacks. **(G)** Analysis of appressorium penetration. Maize leaves infected with SG200 and SG200 *Δlem2* were stained with chlorazol black at 24 hpi to visualize fungal structures. Left panel: Quantification of the different infection stages for each strain. Right panel: Representative Z-axis projections illustrating the quantified categories. Within these images, magenta asterisks indicate the site of appressorium formation and initial penetration, magenta arrows highlight invasive hyphae, and black circles denote clamp cells. In panels (A, C, D, F, G), error bars represent the SD from three independent replicates. Statistical significance was determined using an ordinal logistic mixed model (GLMM) for panels (A, D) and (G), and a Student’s t-test for panels (C) and (F) (ns, not significant; ****, *P* < 0.0001). The total number of infected plants (A) or cells (C, D, F, G) analyzed is indicated above each column.

To characterize the basis of this avirulence, we analyzed fungal colonization by confocal microscopy, which revealed that the *Δlem2* mutant failed to colonize the plant tissue in both genetic backgrounds (Fig. 2B). Given the absence of proliferative hyphae inside the plant, we next analyzed early steps of the pathogenic program to determine which stage of infection is impaired. We first assessed conjugation tube formation in haploid FB1 cells using a pheromone stimulation assay. Exposure to synthetic pheromone revealed no differences in conjugation tube formation between the wild-type and *Δlem2* strains (Fig. 2C); furthermore, subsequent nuclear translocation into the conjugative filament was unaffected in the mutant (Fig. 2D). Since mating on charcoal-containing medium was also unaffected in the *Δlem2* mutant (Fig. 2E), we turned our attention to later stages of pathogenesis using the solopathogenic strain SG200. We began by assessing appressorium formation using the AM1::GFP reporter, which specifically marks filaments undergoing appressorium differentiation. These assays revealed no differences in appressorium formation between the wild-type and the *Δlem2* strain (Fig. 2F). However, quantitative analysis revealed a specific defect in penetration. In the *Δlem2* mutant, 73% of appressoria failed to penetrate, and an additional 13% arrested immediately post-penetration. In contrast, 86.5% of wild-type appressoria successfully penetrated and proliferated within the plant tissue (Fig. 2G). Consistently, the FB1×FB2 *Δlem2* mutants also exhibited defects in appressorium penetration (Fig. S2). Thus, Lem2 is dispensable for appressorium differentiation but strictly required for host penetration.

### Loss of Lem2 leads to a failure in the transcriptomic reprogramming required for nuclear migration

Plant penetration necessitates the translocation of the nucleus from the appressorium into the nascent invasive hypha (59). Because capturing the precise moment of nuclear exit within plant tissue is technically challenging, we utilized the AB33 laboratory strain to study these dynamics. In this strain, the compatible mating type bE2/bW1 genes are under the control of the nitrate-inducible *nar1* promoter (60), enabling the synchronized induction of infective-like filaments in culture and providing a tractable model to monitor nuclear movement in real-time. Given the NE defects observed in the *Δlem2* mutant, we hypothesized that the mechanical forces required for nuclear translocation might compromise or rupture the NE. Contrary to this hypothesis, live-cell imaging revealed that the NE remained intact during the process. However, we observed a striking defect in nuclear exit into the filament. While wild-type nuclei efficiently migrate into the budding hypha, *Δlem2* nuclei moved to the cell pole but failed to traverse the neck, remaining arrested at the yeast-filament junction (Fig. 3A). This blockage occurred in approximately 80% of the cells (Fig. 3B). Notably, the remaining 20% of *Δlem2* nuclei successfully completed migration, indicating that the core machinery for nuclear movement is present and functional in the mutant (Fig. 3B and C). This observation, combined with the lack of NE rupture and the normal organization of actin and microtubules (Fig. S3A, B), suggests that the nuclear migration defect stems from a regulatory rather than a structural or mechanical failure. To identify the transcriptional changes underlying this phenotype, we performed RNA-sequencing (RNA-seq) at 0h and 3.5 h post-induction, coinciding with the onset of nuclear exit in the wild-type strain (Fig. S3C).

**Figure 3.**
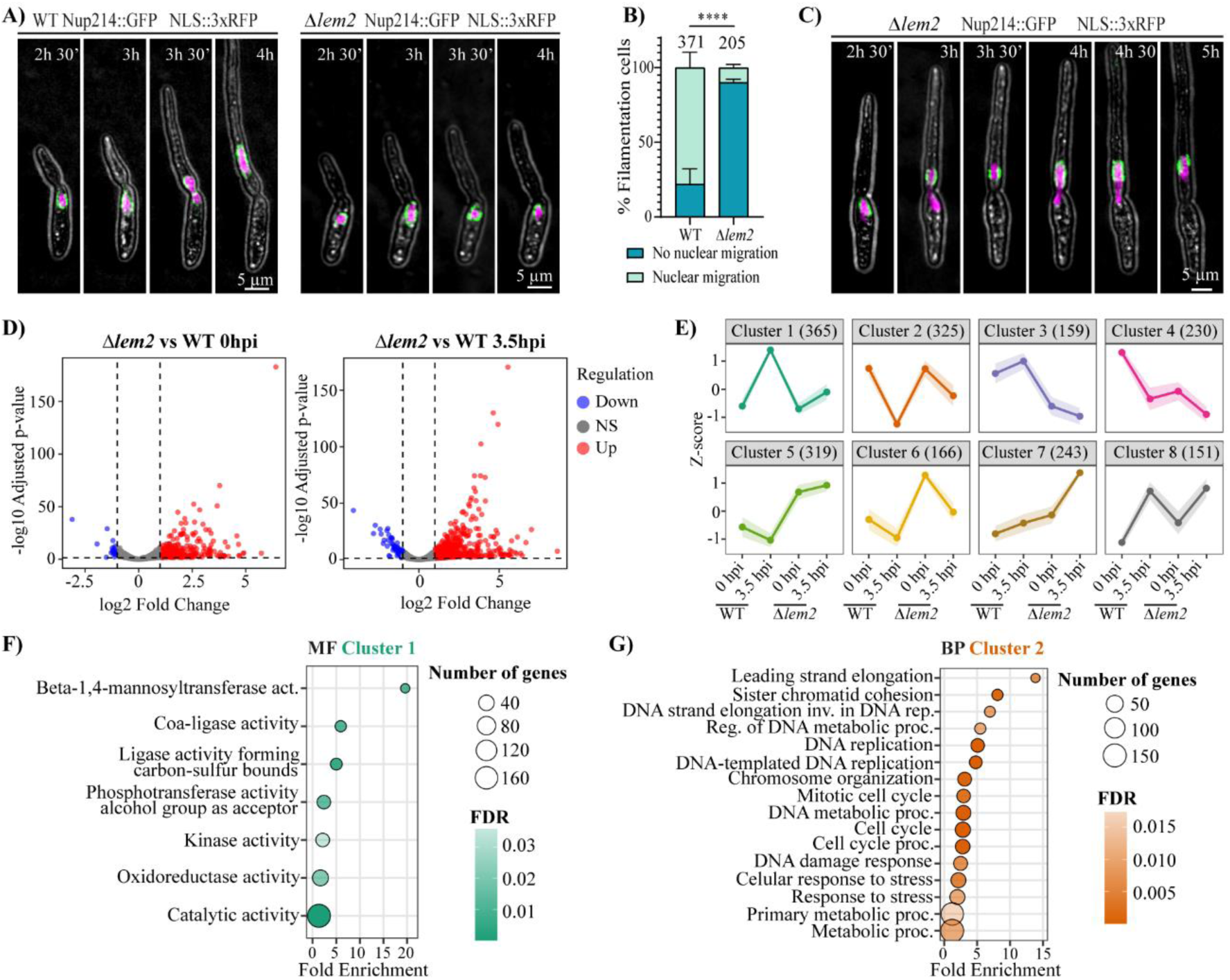
Lem2 is required for proper nuclear migration during filamentation. **(A)** Time-lapse imaging of nuclear migration in WT and *Δlem2* filaments. Consecutive frames illustrate the dynamics of filament elongation and nuclear movement. Strains express Nup214::GFP (green) and NLS::3xRFP (magenta). Numbers above each frame indicate the time in minutes after induction of the filamentation. **(B)** Quantification of nuclear migration determined by DAPI staining. Error bars represent the SD from three independent replicates. **(C)** Representative time-lapse frames of *Δlem2* filaments showing a mutant capable of successful nuclear migration. Numbers above each frame indicate the time in minutes after induction of the filamentation. **(D)** Volcano plots of differentially expressed genes (DEGs). Transcriptomic profiles of *Δlem2* are compared relative to WT at 0 hpi (left) and 3.5 hpi (right). Red dots represent significantly upregulated genes (log2FC > 1), and blue dots represent significantly downregulated genes (log2FC < -1). **(E)** K-means clustering of gene expression profiles. To identify transcriptional patterns, genes were grouped based on Z-scores of normalized counts (VST-transformed) across all conditions. **(F–G)** Functional enrichment analysis. Gene Ontology (GO) terms significantly enriched in Molecular Function (MF) for Cluster 1 **(F)** and in Biological Process (BP) for Cluster 2 **(G)**. The analysis was performed using ShinyGO 0.85 (FDR < 0.05). In (B), significance was determined using the GLMM (****, *P* < 0.0001). The total number of filaments analyzed (*n*) is indicated above each column. All panels feature strains in the AB33 genetic background.

The transcriptomic analysis revealed that the loss of Lem2 results in the deregulation of many genes at both studied time points (813 and 1600 at 0 hpi and 3.5 hpi, respectively). In the *Δlem2* mutant, a large proportion of these differentially expressed genes (DEGs) were significantly upregulated (70.4% and 63.4% at 0 and 3.5 hpi, respectively), consistent with a primary role for Lem2 as a silencing factor (Fig. 3D). Clustering analysis identified two groups of particular interest, clusters 1 and 2, which represent genes that undergo a characteristic transcriptional shift during nuclear exit in wild-type cells, but fail to be properly regulated in the absence of Lem2 (Fig. 3E). Cluster 1 comprises genes that are induced during nuclear exit in the wild-type but fail to reach these induction levels in the *Δlem2* mutant. Gene Ontology (GO) analysis of this cluster revealed an enrichment in terms related to kinase activity pathways (Fig. 3F). This group includes essential regulators of the pathogenic development pathway, as the core members bE1 and Clp1, as well as Kpp6 and Hdp2, two downstream members specifically associated with appressorium penetration (61). Conversely, cluster 2 includes genes that are downregulated during nuclear exit in the wild type but are not efficiently repressed in the *Δlem2* mutant. This cluster is enriched in terms for the cell cycle, including the cyclins 1 and 2 (Clb1, Clb2), almost the entire spindle assembly checkpoint (SAC) pathway, and key regulatory kinases such as ALK1, MPH1, and CDC5. Additionally, the cluster is enriched in terms for DNA replication and the DNA damage response (DDR) featuring the RFC complex (RFC4, RFC5), the Minichromosome Mantenance (MCM) complex components (CDC46, MCM10, MCM2/NimQ), chromatin remodelers (Hat1, Spt16), and specialized factors for replicative stress and homologous recombination at telomeres, such as the cohesin Smc1, the recombinase Rec2, and the Pif1-family helicase Rrm36 (Fig. 3G).

### The MSC domain of Lem2 is essential for telomere anchoring and nuclear migration

The phenotypic and transcriptional defects observed in the *Δlem2* mutant could arise from telomere detachment or other structural roles of Lem2. Interestingly, we found that while tagging the shelterin protein Rap1 with GFP did not interfere with nuclear migration (Fig. 4A and B) or telomere anchoring (Fig. S4A), tagging the distal-most shelterin component, Pot1, resulted in a nuclear migration defect similar to that of the *Δlem2* mutant (Fig. 4A and B). Notably, in Pot1-GFP cells, telomeres remained properly anchored to the NE (Fig. S4A). These results indicate that perturbing the distal-most component of the shelterin complex is sufficient to hinder nuclear translocation, suggesting that telomere stability, beyond simple physical attachment, is a critical requirement for nuclear migration. In the *Δlem2* mutant, telomere stability is likely compromised because, unlike in the wild-type strain, telomeres detach from the nuclear periphery during filamentation (Fig. 4C), a defect that likely contributes to the observed migration failure. To further explore this, we investigated the role of the conserved MSC domain, which in other fungi specifically mediates telomere anchoring (23). We complemented the *Δlem2* mutant with either the full-length protein, the C-terminal region (containing the transmembrane and MSC domains), or the N-terminal region (containing the transmembrane and LEM domains). The MSC domain was sufficient to restore telomere anchoring to the NE (Fig. 4D, E). In contrast, expression of the LEM domain disrupted NE integrity, which precluded the quantification of telomere positioning (Fig. S4B).

**Figure 4.**
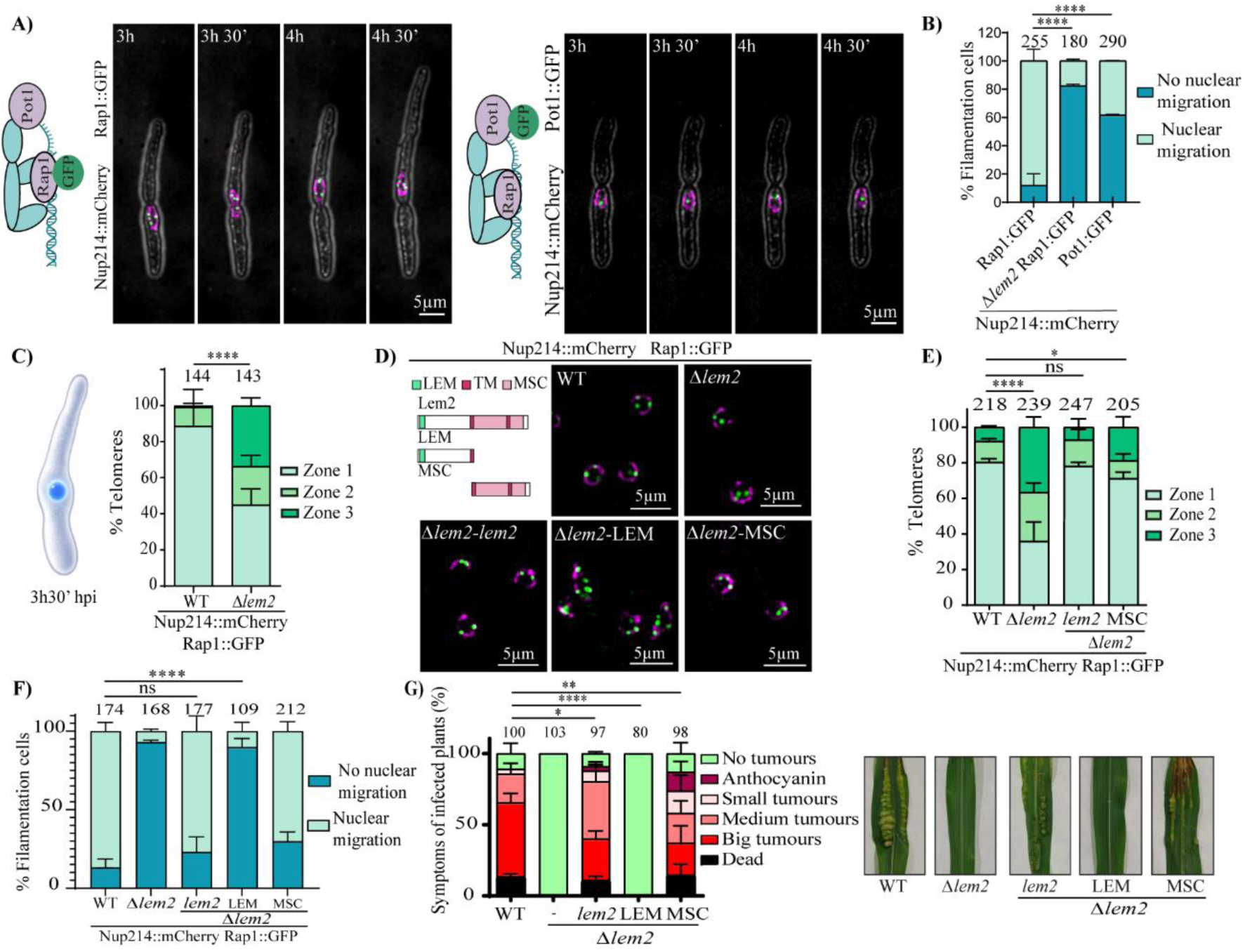
Telomere re-anchoring to the nuclear envelope is essential for nuclear migration. **(A)** Representative time-lapse frames of filaments expressing Nup214::mCherry (magenta) and either Rap1::GFP or Pot1::GFP (green). Numbers above each frame indicate the time in minutes after induction of the filamentation. **(B)** Quantification of nuclear migration in the indicated strains using DAPI staining. **(C)** Quantification of telomere positioning in WT and *Δlem2* during filamentation, just before nuclear exit into the filament. (Left) Schematic representation of the specific stage analyzed. **(D)** Schematic representation of Lem2 domain architecture and representative images of telomere localization in *Δlem2* cells complemented with either the MSC or the LEM domain. TM, transmembrane domain. **(E)** Quantification of telomere distribution relative to the NE for the indicated strains. **(F)** Quantification of nuclear migration in the indicated strains using DAPI staining. **(G)** Pathogenicity assays in maize plants. Left panel: Quantification of disease symptoms at 14 dpi for the indicated strains. Right panel: Representative images of plant infection symptoms. In panels (B, F, G), error bars represent the SD from three independent replicates. Statistical significance was determined using the ordinal logistic mixed model (GLMM) (ns, not significant; ****, *P* < 0.0001). The total number of filaments or plants analyzed (*n*) is indicated above each column. All panels feature strains in the AB33 genetic background, except for panel G, which uses the SG200 background.

Importantly, reintroduction of the MSC domain also rescued nuclear migration during filamentation in the AB33 *Δlem2* background (Fig. 4F), and partially restored both host colonization (Fig. S4C) and overall pathogenicity in the SG2OO solopathogenic background (Fig. 4G). Conversely, the LEM domain failed to complement either nuclear migration or virulence (Fig. 4F and G). Together, these results demonstrate that in *U. maydis*, the MSC domain of Lem2 is essential for telomere anchoring and is required for proper nuclear migration, suggesting a tight functional link between telomere organization at the NE and fungal pathogenesis.

### Telomere anchoring to the nuclear envelope is essential for nuclear migration and host penetration

To determine whether telomere detachment is the primary cause of the nuclear translocation defect in the *Δlem2* mutant, we artificially tethered telomeres to the NE and assessed the impact on both nuclear movement and pathogenesis. We utilized a recruitment system consisting of a GFP-binding protein (GBP) fused to the nucleoporin Nup84 in a *Δlem2* strain expressing Rap1-GFP. This targeted approach successfully re-anchored telomeres to the nuclear pore complexes (Fig. 5A). While this strategy induced telomere declustering in both wild-type and *Δlem2* cells, control experiments showed that declustering did not compromise filamentation or nuclear migration in wild-type cells (80% migration) (Fig. 5B and S5A), validating the system for functional rescue assays. Strikingly, GBP-mediated re-anchoring of telomeres partially rescued both the nuclear migration defect (Fig. 5B and C) and fungal penetration (Fig. 5D and S5B) into plant tissues in *Δlem2* cells. Although telomere re-anchoring restores plant penetration, it is not sufficient to restore macroscopic tumor formation in maize (Fig. 5E and S5C), suggesting additional roles for Lem2 during later stages of infection. These results demonstrate that physical anchoring of telomeres to the NE is a critical requirement for proper nuclear migration and plant penetration.

**Figure 5.**
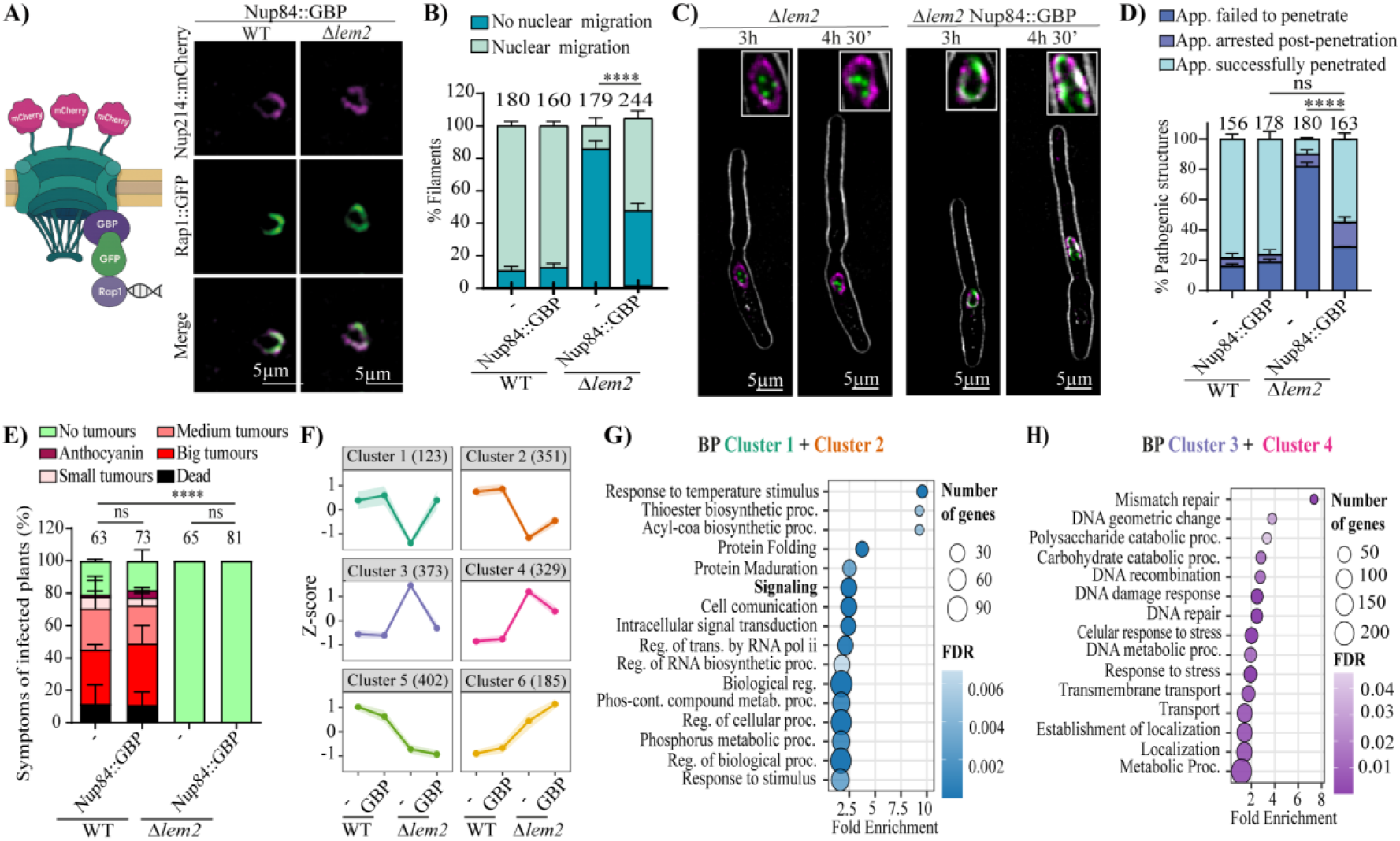
Artificial telomere anchoring rescues nuclear migration and plant penetration in the *Δlem2* mutant. **(A)** Left, schematic representation of the GBP-based recruitment system used to tether telomeres to the nuclear envelope (NE). Right, representative images of WT and *Δlem2* cells expressing Rap1::GFP (green) and Nup214::mCherry (magenta) in the presence of Nup84::GBP to confirm telomere re-anchoring at the NE. **(B)** Quantification of nuclear migration status in the indicated strains at 5 hours post-induction (hpi), determined by DAPI staining. **(C)** Representative time-lapse frames of nuclear migration during filamentation in the indicated strains. Numbers above each frame indicate the time in minutes after induction of the filamentation. **(D)** Analysis of appressorium penetration. Maize leaves infected with the indicated strains were stained with chlorazol black at 24 hpi. **(E)** Disease symptoms in maize plants infected at 14 dpi. The total number of infected plants is indicated above each column. **(F)** K-means clustering of gene expression profiles. Genes were grouped based on their normalized expression levels (counts) across all conditions to identify transcriptional patterns. **(G–H)** Functional enrichment analysis. Gene Ontology (GO) terms significantly enriched among genes in Cluster 1 + Cluster 2 (G) and Cluster 3 + Cluster 4 (H). The analysis was performed using ShinyGO 0.85 (FDR < 0.05). In panels (C, D, E), error bars represent the SD from three independent replicates. Statistical significance was determined using the ordinal logistic mixed model (GLMM) (ns, not significant; ****, P < 0.0001). The total number of filaments (*n*) or plants analyzed is indicated above each column. All panels feature strains in the AB33 genetic background.

Finally, we performed transcriptomic analysis to identify the regulatory pathways complemented by telomere re-anchoring. Re-attachment of telomeres to the NE led to a substantial transcriptional recovery, as evidenced by a significant reduction in the number of DEGs compared to the *Δlem2* mutant (Fig. S5D). To identify the specific genes responsible for the rescue of nuclear migration, we performed a clustering analysis. Clusters 1 and 2 comprised genes downregulated in the *Δlem2* mutant whose expression was restored to near wild-type levels upon telomere re-anchoring (Fig. 5F). These clusters showed an enrichment in signaling pathways (Fig. 5G), specifically the MAPK signaling pathway and core pathogenic regulators (e.g., *clp1, hdp2*) (Fig. S5E). Furthermore, Clusters 3 and 4 identified genes upregulated in the *Δlem2* mutant that were effectively repressed following artificial anchoring, including those involved in DNA damage and repair (e.g., *rec2*, *rrm36*) (Fig. 5H). Notably, while Nup84-tethering successfully resolved this acute DNA damage signature, a subset of upregulated genes related to cell cycle progression and the SAC pathway (such as *clb1* and *mad2*) remained deregulated and were not fully repressed. Together, these findings establish that telomere positioning at the NE acts as a key regulatory signal for the transcriptomic reprogramming and nuclear dynamics essential for fungal infection.

## Discussion

The spatial organization of the genome within the nucleus is a highly regulated process rather than a random occurrence (62). A fundamental hallmark of this architecture is the sequestration of heterochromatin at the nuclear periphery, a specialized microenvironment that provides the necessary epigenetic landscape for transcriptional silencing (23, 63). Beyond its role as a repressive scaffold, the nuclear envelope (NE) has emerged as a sophisticated signaling hub that integrates mechanical and regulatory cues (64). Crucial pathways, including the DNA Damage Response (DDR), MAPK signaling, and cell cycle checkpoints (SAC), are coordinated at this interface to ensure genomic stability and orchestrate complex developmental transitions (65, 66). In this work, we explored the importance of nuclear architecture in fungal phytopathogenesis by characterizing Lem2, the sole LEM-domain protein in the maize pathogen *Ustilago maydis*. We found that Lem2 is indispensable for telomere tethering to the NE, a physical requirement that is essential for nuclear migration and host penetration, thereby linking chromatin spatial positioning directly to fungal virulence.

### Evolutionary conservation and functional specialization of Lem2

In *U. maydis*, Lem2 is critical for NE stability, telomere anchoring, and gene silencing. These roles reflect a fundamental functional conservation across eukaryotes, despite the absence of a classical nuclear lamina in fungi (20). Studies utilizing yeast models, primarily *Saccharomyces cerevisiae* and *Schizosaccharomyces pombe*, have identified functional LEM domain-containing proteins that share a conserved architecture: two transmembrane domains that allow both the N-terminal LEM domain and the C-terminal MSC domain (Man1-Src1p C-terminal domain) to be exposed to the nucleoplasm (21, 22). While both *S. cerevisiae* (Heh1/Src1 and Heh2) and *S. pombe* (Lem2 and Man1) possess two such proteins, *U. maydis* relies on Lem2 as the sole representative of this family, suggesting a consolidation of essential nuclear functions into a single molecular scaffold. In *S. cerevisiae*, Src1/Heh1 is essential for nucleolar integrity and subtelomeric tethering and silencing (67–69), while also participating in nuclear pore complex (NPC) quality control alongside its paralog Heh2 (70, 71). In *S. pombe*, Lem2 and Man1 are important for nuclear envelope structure and involved in heterochromatin tethering and silencing (22–30). Specifically, in Lem2, the LEM domain coordinates centromere clustering (23), whereas the MSC domain, which adopts a winged helix-turn-helix (wHTH) fold, is the primary determinant for telomere localization and silencing through the recruitment of the Snf2/Hdac Repressive Complex (SHREC) (22, 23). This functional specialization is also observed in *U. maydis*, where the MSC domain, but not the LEM domain, is essential for silencing and telomere anchoring. Furthermore, our finding that overexpression of the N-terminal region affects NE integrity is consistent with this region being essential to compensate for the growth arrest of a *lem2Δ lnp1Δ* mutant in *S. pombe,* which is characterized by severe NE defects (29).

Beyond these striking similarities, we identified subtle but significant divergences. Lem2 appears more critical for NE integrity in *U. maydis* than in *S. pombe*. While *Δlem2* mutants in *S. pombe* exhibit NE alterations detectable only by electron microscopy (22), they do not show the prominent nuclear protrusions visible via GFP-NLS (29) that characterize the *U. maydis* mutant. Additionally, Lem2 localization differs between these species; whereas Lem2 is homogeneously distributed across the entire NE in *U. maydis*, it is predominantly concentrated at a single dot colocalizing with the Spindle Pole Body (SPB) in *S. pombe* (22). These differences may be explained by the lack of other LEM-containing proteins in the *U. maydis* genome. In this context, *U. maydis* Lem2 likely consolidates the essential structural and regulatory roles that are otherwise diversified among multiple lamina-associated proteins (LAPs) in other eukaryotic lineages.

### Telomere detachment triggers a surveillance response that restricts nuclear migration and fungal pathogenesis

At the natural ends of linear chromosomes, telomeres must be tightly protected from being recognized as DNA double-strand breaks (DSBs) to prevent the aberrant activation of the DNA damage response (DDR) (72). This essential capping function is primarily mediated by the shelterin complex (73). Loss of this complex leads to telomere uncapping, triggering a profound DDR governed by the ATM and ATR kinase signaling cascades (74). However, DNA repair factors also actively promote telomere maintenance. For instance, in humans, the tumor suppressor BRCA2, a key component of the homologous recombination pathway that loads the RAD51 recombinase at DSB sites, facilitates telomere replication (75–77). Thus, a delicate balance and precise regulation of these repair functions at telomeres are required. Unlike standard yeast models, *U. maydis* serves as an exceptional system for studying DNA repair and telomere function; its recombinational repair machinery shares key features with metazoans, and it remarkably possesses the exact same 6-bp telomeric repeat sequence found in animals (78–80). Indeed, just as in human cells, the *U. maydis* homologs of BRCA2 and RAD51 are essential for telomere maintenance (81). Rad51 binds directly to the shelterin protein Pot1, facilitating telomere preservation. However, in the absence of Pot1, Rad51 and Brh2 (the BRCA2 homolog) promote abnormal recombination at these sites (82). Similarly, just as Pot1 prevents the aberrant activity of recombination factors, the Ku repair complex, which typically binds to DSBs, is essential in *U. maydis* to suppress the unwanted activation of the DDR at telomeres, a scenario that would otherwise cause a constant cell cycle arrest (83). In this context, the stability of telomeric protein complexes is clearly essential for DDR suppression. Although telomere detachment from the nuclear periphery in *S. pombe* does not reportedly induce the DDR (84), our transcriptomic analysis indicates that the loss of telomere anchoring to the NE in *U. maydis* causes DDR induction. We hypothesize that Lem2-mediated anchoring at the nuclear periphery is required to either maintain sheltering component stability or spatially restrict recombination factors like Brh2 and RAD51 (or its highly active homolog Rec2) from inappropriately accessing the telomeres. Indeed, our data shows that the loss of Lem2 triggers a distinct transcriptional signature of replicative stress. The simultaneous upregulation of chromatin remodelers (*hat1*, *spt16*), the specialized telomeric helicase *rrm36*, and components of the replication machinery (*rfc4*, *rfc5*, *cdc46*) suggests a coordinated effort to open and process these telomeric regions. Together with the cohesin smc1 and the recombinase rec2, this expression profile strongly implies a persistent attempt to rescue these unanchored ends via homologous recombination.

Intriguingly, in *U. maydis*, the activation of the DDR, specifically the Atr1 and Chk1 kinases, is actively required for infection, as it orchestrates a transient G2 arrest that permits initial pathogenic development (54, 85–87). Conversely, we observe that the DDR induced by telomere detachment blocks infection at the appressorium penetration stage and halts nuclear migration. One possibility is that this paradox lies in the divergent nature of the DDR triggers. The physiological G2 arrest required for virulence is a highly regulated developmental signal occurring independently of actual DNA damage or canonical sensor complexes, such as MRN and the 9-1-1 clamp (87). In contrast, telomere detachment induced by the absence of Lem2 causes a genuine genomic damage crisis. This triggers a persistent DDR, which locks the cells in a pathological G2 arrest accompanied by the uncoordinated accumulation of cyclins (Clb1, Clb2) and SAC components, such as Mad2. We propose this acute replicative stress overrides proper developmental cell cycle controls and leads to the repression of the pathogenic program, as evidenced by the failure to fully induce core developmental regulators (*bE1*, *clp1*) and their downstream effectors (*kpp6*, *hdp2*) in the *Δlem2* mutant. Crucially, artificial re-anchoring of telomeres to the Nup84 protein at the NE is sufficient to reduce DDR induction, restore the MAPK signaling pathway, and permit initial penetration. This suggests that the nuclear periphery acts as a protective microenvironment essential for telomere maintenance. While this rescue could simply be the result of regaining general peripheral localization, the specific role of Nup84 in yeast DNA repair makes an alternative hypothesis highly attractive. In yeast, DSBs and eroded telomeres actively relocate to nuclear pore complexes (NPCs) to be properly repaired (88–90). Specifically, the Nup84 subcomplex is involved in DSB recruitment to the NPCs and is essential for telomere localization to the nuclear envelope and efficient DNA double strand break repair at subtelomeric regions (90, 36). Thus, it is tempting to propose a model in which telomeres must reside at the nuclear periphery, anchored by Lem2, to maintain constant proximity to nuclear pores, ensuring their structural and replicative maintenance. When telomeres are detached following *lem2* deletion, they incur damage, whether through shelterin complex instability, increased distance from nuclear pores, or both. This damage induces a constitutive DDR, fundamentally different from the transient, damage-independent DDR essential for infection, ultimately repressing the pathogenic program. Interestingly, while Nup84-mediated tethering successfully suppresses this DDR (Clusters 3 and 4) and reactivates the MAPK pathway (Clusters 1 and 2) to permit initial host invasion, it does not restore macroscopic tumor formation. We hypothesize that this outcome uncouples two distinct regulatory functions of the nuclear periphery. Proximity to NPCs provides the structural microenvironment needed to prevent telomeric damage and turn off the acute DDR, thereby releasing the block on appressorium development. However, Nup84 likely cannot substitute for the specific structural and topological dynamics provided by Lem2 during the demanding structural changes of appressorium differentiation and nuclear migration. This chronic topological defect likely prevents the proper resetting of the cell cycle machinery, ultimately precluding the coordinated mitotic divisions required for sustained tumor proliferation inside the host.

In this work, we show how physical telomere detachment triggers an unresolvable DNA damage response that aborts development, directly linking spatial genome organization to cellular fate. Given the strict evolutionary conservation of LEM-domain proteins, it will be of great interest to investigate whether this spatial requirement for telomere stability and virulence operates in other fungal pathogens. Furthermore, this protective mechanism likely extends beyond fungi; indeed, peripheral instability is a recognized hallmark of metazoan aging and cancer. Ultimately, alongside unveiling novel targets for antifungal design, our study highlights how eukaryotes integrate nuclear architecture, telomere homeostasis, and cell cycle checkpoints to govern complex developmental transitions.

## Materials and Methods

### Strains and growth conditions

*U. maydis* strains used in this study are derived from FB1, FB2, SG200, and AB33 backgrounds and are listed in Table S1 (SI Appendix). Cells were grown as previously described (91). Strains were generated by transforming *U. maydis* protoplasts with the required genetic constructs, as previously described (92). Correct integration into the target loci was validated in all cases by PCR and Southern blot analysis. Detailed procedures for strain construction and growth conditions are provided in the SI Appendix.

### Plasmid Construction

For plasmid assembly, we utilized NEBuilder HiFi DNA Assembly (New England BioLabs) or standard molecular cloning strategies (93) using the plasmids pGEM-T Easy (Promega), pBSK and p123 (94). Specific primers, plasmids and cloning strategies are summarized in S1 appendix, Table S2.

### Plant infection assays

To evaluate the virulence of the different *U. maydis* strains, pathogenicity assays were conducted using *Zea mays* (cv. Early Golden Bantam). Detailed procedures for plant infection assays are provided in SI Appendix.

### Sequence Alignment and Phylogenetic Analysis

The *U. maydis* Lem2 protein (UMAG_00208) was identified via BlastP using fungal, and yeast homologs. Sequence alignments and phylogenetic trees were inferred using MAFFT (v.7) with 1,000 bootstrap replicates to ensure branch reliability. Detailed bioinformatic parameters and the list of species included in the analysis are provided in the SI Appendix.

### Sample preparation for microscopy

Live-cell imaging was performed using Spinning Disk or DeltaVision systems. To assess telomere distribution, the nuclear area in each optical section was divided into three concentric zones of equal area (58), representing the nuclear periphery (Zone I), the intermediate region (Zone II), and the nuclear interior (Zone III). The distance between telomeric foci and the nuclear envelope was quantified using ImageJ/Fiji. *U. maydis* progression within maize tissue was monitored via WGA-Alexa Fluor 488 and propidium iodide staining as described (95). Early fungal structures were visualized using Chlorazole Black (96). For actin and tubulin immunofluorescence, cells were processed as previously described (93), except that the methanol treatment was omitted. Specific technical settings and microscope features are detailed in the SI Appendix.

### Induction of *U. maydis* Infective Structures

To assess conjugation tube formation, FB1 and FB1 *Δlem2* strains were induced by the addition of 1 µL of a2 pheromone (2.5 mg/mL), followed by incubation at 25°C for 5 hours with constant agitation. As a control, separate aliquots were incubated with 1 µL of DMSO. Conjugation tube formation was observed via differential interference contrast (DIC) and fluorescence microscopy using a DeltaVision system.

In vitro appressorium induction was performed on Parafilm M surfaces as previously described (52, 97). Cell cultures in 2% YEPSL were sprayed using an EcoSpray atomizer with 100 µM 16-hydroxyhexadecanoic acid. At 18 hpi, samples were stained with Calcofluor White (CFW, 1 µg/mL). Appressoria were quantified as the percentage of CFW-stained structures relative to GFP-expressing filaments.

### RNA extraction and RNA-Seq analysis

Total RNA was extracted from *U. maydis* AB33 background strains at 0 h and 3.5 h post-induction of filamentation using TRIzol (Invitrogen) and the Direct-zol RNA Miniprep Plus Kit (Zymo Research). Three biological replicates were processed for each condition. Library preparation and paired-end sequencing (PE100) were performed by BGI (Hong Kong) using the DNBseq platform. Differential expression analysis was performed using DESeq2, considering genes with an adjusted p-value <0.05 and |log2FC| ≥0.5 as significant. For these genes, variance-stabilizing transformed (VST) counts derived from the DESeq2 model were averaged by condition, scaled by gene (z-score), and clustered using k-means to identify expression patterns. Details on bioinformatics analysis are provided in Supplementary Methods.

### Data, Materials, and Software Availability

The data that support the findings of this study are openly available in (repository name and URL will be available after acceptance), reference number (reference number will be available after acceptance), and in the supplementary material of this article.

## Acknowledgments

We would like to thank the Genetics Department for their useful discussions and comments. Victor Manuel Carranco, Sandra Romero and Ismael Fernández Portillo for the technical assistant. This research was supported by MCIN/AEI/10.13039/501100011033/ grant number PID2019-110477GB-I00 to JII, by MCIN/AEI/10.13039/501100011033/ and FEDER, UE grant number PID2022-142783NB-I00 to RRB and JII , by Junta de Andalucía grant number UPO-1381372 to JII and by Pablo de Olavide University grant number PPI2304 to RRB.

## Author contributions

E.S-M, J.I.I and R.R.B planned and designed the research. E.S-M, M.V-G and R.R.B generate strains, performed the experiments, and analyzed the data. E.S-M and B.N and R.R.B performed and analyzed RNA-seq experiments. E.S-M and R.R.B wrote the original manuscript with input from all coauthors.

## References

1. B. van Steensel, A. S. Belmont, Lamina-Associated Domains: Links with Chromosome Architecture, Heterochromatin, and Gene Repression. Cell 169, 780–791 (2017).

2. A. Taddei, S. M. Gasser, Structure and Function in the Budding Yeast Nucleus. Genetics 192, 107 (2012).

3. M. Amendola, B. V. Steensel, Mechanisms and dynamics of nuclear lamina–genome interactions. Current Opinion in Cell Biology 28, 61–68 (2014).

4. A. Mattout, D. S. Cabianca, S. M. Gasser, Chromatin states and nuclear organization in development - a view from the nuclear lamina. Genome Biology 16, 174- (2015).

5. J. C. Harr, A. Gonzalez-Sandoval, S. M. Gasser, Histones and histone modifications in perinuclear chromatin anchoring: from yeast to man. EMBO reports 17, 139–155 (2016).

6. N. S. Alagna, T. I. Thomas, K. L. Wilson, K. L. Reddy, Choreography of lamina-associated domains: structure meets dynamics. FEBS Letters 597, 2806–2822 (2023).

7. D. A. Agard, J. W. Sedat, Three-dimensional architecture of a polytene nucleus. Nature 1983 302:5910 302, 676–681 (1983).

8. H. Funabiki, L. Hagan, S. Uzawa, M. Yanagida, Cell cycle-dependent specific positioning and clustering of centromeres and telomeres in fission yeast. Journal of Cell Biology 121, 961–976 (1993).

9. I. Gonzalez-Suarez, et al., Novel roles for A-type lamins in telomere biology and the DNA damage response pathway. EMBO Journal 28, 2414–2427 (2009).

10. M. Hochstrasser, D. Mathog, Y. Gruenbaum, H. Saumweber, J. W. Sedat, Spatial organization of chromosomes in the salivary gland nuclei of Drosophila melanogaster. Journal of Cell Biology 102, 112–123 (1986).

11. A. Ottaviani, et al., Identification of a perinuclear positioning element in human subtelomeres that requires A-type lamins and CTCF. The EMBO Journal 28, 2428 (2009).

12. B. D. Towbin, P. Meister, S. M. Gasser, The nuclear envelope — a scaffold for silencing? Current Opinion in Genetics & Development 19, 180–186 (2009).

13. Y. Fang, D. L. Spector, Centromere Positioning and Dynamics in Living Arabidopsis Plants. 10.1091/mbc.e05-08-0706 16, 5710–5718 (2005).

14. K. L. Wilson, J. M. Berk, The nuclear envelope at a glance. Journal of Cell Science 123, 1973–1978 (2010).

15. N. Briand, P. Collas, Lamina-associated domains: peripheral matters and internal affairs. Genome Biology 2020 21:1 21, 85- (2020).

16. N. Wagner, G. Krohne, LEM-Domain Proteins: New Insights into Lamin-Interacting Proteins. International Review of Cytology 261, 1–46 (2007).

17. S. M. Schreiner, P. K. Koo, Y. Zhao, S. G. J. Mochrie, M. C. King, The tethering of chromatin to the nuclear envelope supports nuclear mechanics. Nature Communications 2015 6:1 6, 7159- (2015).

18. P. Malik, et al., Cell-specific and lamin-dependent targeting of novel transmembrane proteins in the nuclear envelope. Cellular and Molecular Life Sciences 2010 67:8 67, 1353–1369 (2010).

19. Y. Guo, Y. Kim, T. Shimi, R. D. Goldman, Y. Zheng, Concentration-dependent lamin assembly and its roles in the localization of other nuclear proteins. 10.1091/mbc.e13-11-0644 25, 1287–1297 (2014).

20. L. Koreny, M. C. Field, Ancient Eukaryotic Origin and Evolutionary Plasticity of Nuclear Lamina. Genome Biology and Evolution 8, 2663–2671 (2016).

21. B. J. Mans, V. Anantharaman, L. Aravind, E. V. Koonin, Comparative genomics, evolution and origins of the nuclear envelope and nuclear pore complex. Cell Cycle 3, 1625–1650 (2004).

22. Y. Gonzalez, A. Saito, S. Sazer, Fission yeast Lem2 and Man1 perform fundamental functions of the animal cell nuclear lamina. Nucleus 3, 60–76 (2012).

23. R. R. Barrales, M. Forn, P. R. Georgescu, Z. Sarkadi, S. Braun, Control of heterochromatin localization and silencing by the nuclear membrane protein Lem2. Genes & Development 30, 133–148 (2016).

24. S. Banday, Z. Farooq, R. Rashid, E. Abdullah, M. Altaf, Role of inner nuclear membrane protein complex Lem2-Nur1 in heterochromatic gene silencing. Journal of Biological Chemistry 291, 20021–20029 (2016).

25. Y. Tange, et al., Inner nuclear membrane protein Lem2 augments heterochromatin formation in response to nutritional conditions. Genes to Cells 21, 812–832 (2016).

26. S. Borah, et al., Heh2/Man1 may be an evolutionarily conserved sensor of NPC assembly state. 10.1091/mbc.E20-09-0584 32, 1359–1373 (2021).

27. M. Gu, et al., LEM2 recruits CHMP7 for ESCRT-mediated nuclear envelope closure in fission yeast and human cells. Proceedings of the National Academy of Sciences of the United States of America 114, E2166–E2175 (2017).

28. K. Kume, H. Cantwell, A. Burrell, P. Nurse, Nuclear membrane protein Lem2 regulates nuclear size through membrane flow. Nature Communications 2019 10:1 10, 1871- (2019).

29. Y. Hirano, et al., Lem2 and Lnp1 maintain the membrane boundary between the nuclear envelope and endoplasmic reticulum. Communications Biology 2020 3:1 3, 276-(2020).

30. J. M. Varberg, J. R. Unruh, A. J. Bestul, A. A. Khan, S. L. Jaspersen, Quantitative analysis of nuclear pore complex organization in Schizosaccharomyces pombe. Life Science Alliance 5 (2022).

31. C. Lemaître, et al., Nuclear position dictates DNA repair pathway choice. Genes & Development 28, 2450–2463 (2014).

32. S. Gonzalo, DNA damage and lamins. Advances in Experimental Medicine and Biology 773, 377–399 (2014).

33. L. Etourneaud, et al., Lamin B1 sequesters 53BP1 to control its recruitment to DNA damage. Science Advances 7, 3799–3826 (2021).

34. S. jung Kim, S. H. Park, K. Myung, K. young Lee, Lamin A/C facilitates DNA damage response by modulating ATM signaling and homologous recombination pathways. Animal Cells and Systems 28, 401–416 (2024).

35. B. Moser, J. Basílio, J. Gotzmann, A. Brachner, R. Foisner, Comparative Interactome Analysis of Emerin, MAN1 and LEM2 Reveals a Unique Role for LEM2 in Nucleotide Excision Repair. Cells 2020, Vol. 9, 9 (2020).

36. P. Therizols, et al., Telomere tethering at the nuclear periphery is essential for efficient DNA double strand break repair in subtelomeric region. Journal of Cell Biology 172, 189–199 (2006).

37. N. Agmon, B. Liefshitz, C. Zimmer, E. Fabre, M. Kupiec, Effect of nuclear architecture on the efficiency of double-strand break repair. Nature Cell Biology *2013* 15:*6* 15, 694–699 (2013).

38. B. Lee, T. H. Lee, J. Shim, Emerin suppresses Notch signaling by restricting the Notch intracellular domain to the nuclear membrane. Biochimica et Biophysica Acta (BBA) - Molecular Cell Research 1864, 303–313 (2017).

39. E. Markiewicz, et al., The inner nuclear membrane protein Emerin regulates β-catenin activity by restricting its accumulation in the nucleus. The EMBO Journal 25, 3275 (2006).

40. G. Melcon, et al., Loss of emerin at the nuclear envelope disrupts the Rb1/E2F and MyoD pathways during muscle regeneration. Human Molecular Genetics 15, 637–651 (2006).

41. D. M. Chambers, et al., LEM domain–containing protein 3 antagonizes TGF–SMAD2/3 signaling in a stiffness-dependent manner in both the nucleus and cytosol. Journal of Biological Chemistry 293, 15867–15886 (2018).

42. F. Lin, J. M. Morrison, W. Wu, H. J. Worman, MAN1, an integral protein of the inner nuclear membrane, binds Smad2 and Smad3 and antagonizes transforming growth factor-β signaling. Human Molecular Genetics 14, 437–445 (2005).

43. D. Pan, et al., The Integral Inner Nuclear Membrane Protein MAN1 Physically Interacts with the R-Smad Proteins to Repress Signaling by the Transforming Growth Factor-β Superfamily of Cytokines. Journal of Biological Chemistry 280, 15992–16001 (2005).

44. M. D. Huber, T. Guan, L. Gerace, Overlapping Functions of Nuclear Envelope Proteins NET25 (Lem2) and Emerin in Regulation of Extracellular Signal-Regulated Kinase Signaling in Myoblast Differentiation. Molecular and Cellular Biology 29, 5718 (2009).

45. R. Dean, et al., The Top 10 fungal pathogens in molecular plant pathology. Molecular Plant Pathology 13, 414–430 (2012).

46. M. C. Fisher, et al., Emerging fungal threats to animal, plant and ecosystem health. Nature 484, 186–194 (2012).

47. F. Banuett, Genetics of Ustilago maydis, a fungal pathogen that induces tumors in maize. Annual Review of Genetics 29, 179–208 (1995).

48. R. Kahmann, G. Steinberg, C. Basse, M. Feldbrügge, J. Kämper, Ustilago maydis, the Causative Agent of Corn Smut Disease. Fungal Pathology 347–371 (2000). 10.1007/978-94-015-9546-9_12.

49. T. García-Muse, G. Steinberg, J. Pérez-Martín, Pheromone-Induced G2 Arrest in the Phytopathogenic Fungus Ustilago maydis. Eukaryotic Cell 2, 494 (2003).

50. J. Pérez-Martín, S. Castillo-Lluva, Connections between polar growth and cell cycle arrest during the induction of the virulence program in the phytopathogenic fungus Ustilago maydis. Plant Signaling and Behavior 3, 480–481 (2008).

51. K. M. Snetselaar, C. W. Mims, Light and electron microscopy of Ustilago maydis hyphae in maize. Mycological Research 98, 347–355 (1994).

52. A. Mendoza-Mendoza, et al., Physical-chemical plant-derived signals induce differentiation in Ustilago maydis. Molecular Microbiology 71, 895–911 (2009).

53. K. M. Snetselaar, C. W. Mims, Sporidial Fusion and Infection of Maize Seedlings by the Smut Fungus Ustilago Maydis. Mycologia 84, 193–203 (1992).

54. S. Castanheira, N. Mielnichuk, J. Pérez-Martín, Programmed cell cycle arrest is required for infection of corn plants by the fungus Ustilago maydis. Development 141, 4817–4826 (2014).

55. F. Banuett, I. Herskowitz, Discrete developmental stages during teliospore formation in the corn smut fungus, Ustilago maydis. Development 122, 2965–2976 (1996).

56. D. Lanver, et al., Ustilago maydis effectors and their impact on virulence. Nat Rev Microbiol 15, 409–421 (2017).

57. Y. Hiraoka, et al., Inner nuclear membrane protein Ima1 is dispensable for intranuclear positioning of centromeres. Genes to Cells 16, 1000–1011 (2011).

58. F. Hediger, A. Taddei, F. R. Neumann, S. M. Gasser, Methods for Visualizing Chromatin Dynamics in Living Yeast. Methods in Enzymology 375, 345–365 (2004).

59. J. Kämper, et al., Insights from the genome of the biotrophic fungal plant pathogen Ustilago maydis. Nature *2006* 444:7115 444, 97–101 (2006).

60. A. Brachmann, G. Weinzierl, J. Kämper, R. Kahmann, Identification of genes in the bW/bE regulatory cascade in Ustilago maydis. Molecular Microbiology 42, 1047–1063 (2001).

61. K. Heimel, M. Scherer, D. Schuler, J. Kämper, The Ustilago maydis Clp1 Protein Orchestrates Pheromone and b-Dependent Signaling Pathways to Coordinate the Cell Cycle and Pathogenic Development. The Plant Cell 22, 2908 (2010).

62. W. A. Bickmore, B. V. Steensel, Genome Architecture: Domain Organization of Interphase Chromosomes. Cell 152, 1270–1284 (2013).

63. A. Akhtar, S. M. Gasser, The nuclear envelope and transcriptional control. Nature Reviews Genetics 2007 8:7 8, 507–517 (2007).

64. A. Buchwalter, J. M. Kaneshiro, M. W. Hetzer, Coaching from the sidelines: the nuclear periphery in genome regulation. Nature reviews. Genetics 20, 39 (2019).

65. H. J. Worman, G. G. Gundersen, Here come the SUNs: a nucleocytoskeletal missing link. Trends in Cell Biology 16, 67–69 (2006).

66. K. Mekhail, D. Moazed, The nuclear envelope in genome organization, expression and stability. Nature Reviews Molecular Cell Biology 2010 11:5 11, 317–328 (2010).

67. K. Mekhail, J. Seebacher, S. P. Gygi, D. Moazed, Role for perinuclear chromosome tethering in maintenance of genome stability. Nature 456, 667 (2008).

68. S. E. Grund, et al., The inner nuclear membrane protein Src1 associates with subtelomeric genes and alters their regulated gene expression. The Journal of cell biology 182, 897–910 (2008).

69. J. N. Y. Chan, et al., Perinuclear cohibin complexes maintain replicative life span via roles at distinct silent chromatin domains. Developmental Cell 20, 867–879 (2011).

70. B. M. Webster, P. Colombi, J. Jäger, C. P. Lusk, Surveillance of nuclear pore complex assembly by ESCRT-III/Vps4. Cell 159, 388–401 (2014).

71. B. M. Webster, et al., Chm7 and Heh1 collaborate to link nuclear pore complex quality control with nuclear envelope sealing. The EMBO journal 35, 2447–2467 (2016).

72. T. de Lange, Shelterin-Mediated Telomere Protection. Annu Rev Genet 52, 223–247 (2018).

73. T. de Lange, Shelterin: the protein complex that shapes and safeguards human telomeres. Genes Dev 19, 2100–2110 (2005).

74. J. Maciejowski, T. de Lange, Telomeres in cancer: tumour suppression and genome instability. Nat Rev Mol Cell Biol 18, 175–186 (2017).

75. S. Badie, et al., BRCA2 Acts as RAD51 Loader to Facilitate Telomere Replication and Capping. Nat Struct Mol Biol 17, 1461–1469 (2010).

76. M. Tarsounas, et al., Telomere maintenance requires the RAD51D recombination/repair protein. Cell 117, 337–347 (2004).

77. I. Jaco, et al., Role of Mammalian Rad54 in Telomere Length Maintenance. Mol Cell Biol 23, 5572–5580 (2003).

78. W. K. Holloman, J. Schirawski, R. Holliday, The homologous recombination system of Ustilago maydis. Fungal Genet Biol 45 **Suppl 1**, S31–39 (2008).

79. G. Swapna, E. Y. Yu, N. F. Lue, Single telomere length analysis in Ustilago maydis, a high-resolution tool for examining fungal telomere length distribution and C-strand 5’-end processing. Microb Cell 5, 393–403 (2018).

80. P. A. Guzmán, J. G. Sánchez, Characterization of telomeric regions from Ustilago maydis. Microbiology (Reading) 140 **( Pt** **3**), 551–557 (1994).

81. E. Y. Yu, M. Kojic, W. K. Holloman, N. F. Lue, Brh2 and Rad51 promote telomere maintenance in *Ustilago maydis*, a new model system of DNA repair proteins at telomeres. DNA Repair 12, 472–479 (2013).

82. S. Zahid, S. Aloe, J. H. Sutherland, W. K. Holloman, N. F. Lue, Ustilago maydis telomere protein Pot1 harbors an extra N-terminal OB fold and regulates homology-directed DNA repair factors in a dichotomous and context-dependent manner. PLoS Genet 18, e1010182 (2022).

83. C. de Sena-Tomás, et al., Fungal Ku prevents permanent cell cycle arrest by suppressing DNA damage signaling at telomeres. Nucleic Acids Res 43, 2138–2151 (2015).

84. Y. Chikashige, et al., Membrane proteins Bqt3 and -4 anchor telomeres to the nuclear envelope to ensure chromosomal bouquet formation. J Cell Biol 187, 413–427 (2009).

85. N. Mielnichuk, C. Sgarlata, J. Pérez-Martín, A role for the DNA-damage checkpoint kinase Chk1 in the virulence program of the fungus Ustilago maydis. J Cell Sci 122, 4130–4140 (2009).

86. C. de Sena-Tomás, A. Fernández-Álvarez, W. K. Holloman, J. Pérez-Martín, The DNA Damage Response Signaling Cascade Regulates Proliferation of the Phytopathogenic Fungus Ustilago maydis in Planta[W]. Plant Cell 23, 1654–1665 (2011).

87. M. Tenorio-Gómez, C. de Sena-Tomás, J. Pérez-Martín, MRN- and 9-1-1-Independent Activation of the ATR-Chk1 Pathway during the Induction of the Virulence Program in the Phytopathogen Ustilago maydis. PLOS ONE 10, e0137192 (2015).

88. B. Khadaroo, et al., The DNA damage response at eroded telomeres and tethering to the nuclear pore complex. Nat Cell Biol 11, 980–987 (2009).

89. D. Churikov, et al., SUMO-Dependent Relocalization of Eroded Telomeres to Nuclear Pore Complexes Controls Telomere Recombination. Cell Rep 15, 1242–1253 (2016).

90. S. Nagai, et al., Functional targeting of DNA damage to a nuclear pore-associated SUMO-dependent ubiquitin ligase. Science 322, 597–602 (2008).

91. B. Gillissen, et al., A two-component regulatory system for self/non-self recognition in Ustilago maydis. Cell 68, 647–657 (1992).

92. K. Bösch, et al., Genetic Manipulation of the Plant Pathogen Ustilago maydis to Study Fungal Biology and Plant Microbe Interactions. Journal of visualized experiments : JoVE 2016 (2016).

93. Sambrook, J., Fritsch, E. R., & Maniatis, T. (1989). Molecular Cloning A Laboratory Manual (2nd ed.). Cold Spring Harbor, NY Cold Spring Harbor Laboratory Press. - References - Scientific Research Publishing.

94. C. Aichinger, et al., Identification of plant-regulated genes in Ustilago maydis by enhancer-trapping mutagenesis. Molecular genetics and genomics : MGG 270, 303–314 (2003).

95. A. Redkar, E. Jaeger, G. Doehlemann, Visualization of Growth and Morphology of Fungal Hyphae in planta Using WGA-AF488 and Propidium Iodide Co-staining. (2018). 10.21769/BioProtoc.2942.

96. A. Brachmann, J. Schirawski, P. Müller, R. Kahmann, An unusual MAP kinase is required for efficient penetration of the plant surface by Ustilago maydis. The EMBO Journal *2003* 22:*9* 22, 2199–2210 (2003).

97. D. Lanver, A. Mendoza-Mendoza, A. Brachmann, R. Kahmann, Sho1 and Msb2-Related Proteins Regulate Appressorium Development in the Smut Fungus Ustilago maydis. The Plant Cell 22, 2085–2101 (2010).

