## Supplementary material for "Perinuclear anchoring of telomeres enables plant infection by *Ustilago maydis*": SI appendix

**Methods S1**

**Molecular Cloning and Strain Construction**

DNA cloning and plasmid maintenance were carried out using *Escherichia coli* DH5α following standard procedures (1). For plasmid assembly, we utilized NEBuilder HiFi DNA Assembly (New England BioLabs) or standard molecular cloning strategies (1) using the plasmids pGEM-T Easy (Promega), pBSK and p123 (2). *U. maydis* strains were cultured as previously described (3, 4). Genetic manipulation of *U. maydis*, including genomic DNA isolation and protoplast transformation, was performed according to a modified protocol (5). Specific U. maydis gene numbers for all deletion and tagging constructs are listed in Table S3 (SI Appendix).

To delete *lem2* in *U. maydis* (UMAG_00208), the 5' and 3' flanking regions of the open reading frame were amplified by PCR from genomic DNA using the primer pairs Um00208KO5-1/Um00208KO5-2 and Um00208KO3-1/Um00208KO3-2, respectively (Table S2). These fragments were ligated to a 1.4-kb nourseothricin (NatR) resistance cassette (4) following digestion of both components with SfiI. The ligation products were subsequently cloned into the pGEM-T Easy vector (Promega) to generate the plasmid pGEMT-del-*lem2*::*NatR*. This plasmid served as a template to amplify a linear deletion cassette using primers Um00208KO5-1 and Um00208KO3-2. The resulting PCR product was integrated into the *U. maydis* genome via homologous recombination.

For complementation assays, the plasmid p123 (2) was utilized. This vector contains the constitutive *otef* promoter, a GFP reporter, and a *nos* terminator, along with a carboxin resistance marker (*cbxR*). The resistance marker consists of a mutated version of the *U. maydis* succinate dehydrogenase gene (*sdh2*; UMAG_00844), which allows for targeted integration into the *ip* locus through homologous recombination and selection on carboxin-supplemented media (6). To generate complementation constructs, the GFP sequence in p123 was replaced by the corresponding sequences of the full-length *lem2* ORF or its truncated versions (MSC and LEM domains, both including the endogenous transmembrane domain) amplified from genomic DNA using specific primer pairs (Table S2). This resulted in their expression under the control of the constitutive *otef* promoter following ligation with T4 DNA ligase (New England Biolabs). The resulting plasmids p123-*lem2*, p123-*MSC* and p123-*LEM* were linearized with SspI and integrated into the *ip* locus via homologous recombination.

To generate the Lem2::GFP tagging construct, a strategy based on in vivo recombination in *Saccharomyces cerevisiae* was employed. Two genomic fragments, corresponding to the upstream and downstream regions of the *lem2* stop codon, were amplified by PCR using primers containing homology arms for the pRS426 vector (Table S2). Additionally, a DNA fragment containing the GFP sequence and the hygromycin resistance cassette (*HygR*) was amplified. The pRS426 backbone was linearized by digestion with EcoRI. The linearized vector, the flanking genomic regions, and the GFP-*HygR* cassette were co-transformed into *S. cerevisiae* to allow for plasmid assembly. The resulting plasmid, named pRS426-Um00208:GFP:*HygR*, was subsequently isolated and used as a template to amplify the complete tagging cassette for integration into *U. maydis* strains via homologous recombination.

To visualize nuclear localization, we utilized the plasmid pC_NLS::3xRFP (7), which carries a 3xRFP tag and a carboxin resistance marker (*CbxR*). The plasmid was linearized with *SspI* and integrated into the *ip* locus of the corresponding *U. maydis* strains via homologous recombination. To visualize the endogenous localization of Nup214 (UMAG_01089), we used the plasmid pN214G (8), as a template to amplify a tagging cassette containing a GFP tag and a hygromycin resistance marker (*HygR*) using specific primers (Table S2). The resulting PCR product was integrated into the native *nup214* locus of *U. maydis* strains via homologous recombination.

To visualize Nup214, Rap1, Mis12, Grc1, Pot1, and Nup84, C-terminal tagging was performed at their respective native loci. All tagging plasmids were generated by assembling the required PCR fragments into a BamHI-linearized pBSK vector using NEBuilder HiFi DNA Assembly (New England BioLabs). For each gene of interest (GOI), the 5’ flanking region (KO5), encompassing the C-terminal part of the ORF without the stop codon, and the 3’ flanking region (KO3), corresponding to the downstream region, were amplified from genomic DNA using Q5® High-Fidelity DNA Polymerase. For Rap1, Mis12, Grc1, and Pot1, the assembly included an insert containing GFP fused to a resistance cassette (*HygR* or *GenR*). Similarly, for Nup214, an mCherry-*GenR* fusion fragment was utilized. Regarding the *nup84*::GBP construct, the assembly integrated the *nup84* flanking regions with the GFP-Binding Protein (GBP) sequence, the *nos* terminator, and the carboxin resistance marker (C*bxR*). In all cases, the final tagging cassettes were amplified from the resulting plasmids using the outermost primers and integrated into *U. maydis* strains via homologous recombination.

#### Growth conditions

*U. maydis* strains (listed in Table S1) were cultured in YEPSL medium (0.4% bactopeptone, 1% yeast extract, and 0.4% sucrose) at 28°C (3), unless otherwise specified. For most experimental procedures, cultures were grown to exponential phase and subsequently diluted in fresh medium to synchronize growth before further incubation. For genomic DNA extraction, this second incubation step was omitted, and cultures were harvested directly upon reaching the initial exponential phase.

To trigger filamentous growth, *U. maydis* AB33 derivatives were induced by shifting cells of an exponentially growing culture ($OD_{600}$ of 0.4–0.5) from YEPSL medium to nitrate minimal medium (MM-NO3) supplemented with 1% glucose. Cells were incubated at 28°C with constant shaking at 180 rpm.

To assess mating, cells were grown in YEPSL medium to exponential phase (OD600 0.6–0.8), harvested by centrifugation (850 g for 4 min at room temperature), and washed twice with sterile distilled water. Compatible mating strains (FB1 and FB2) were mixed in a 1:1 ratio, and the final density was adjusted to an $OD_{600}$ of 0.8. Thereafter, a 5 µL aliquot of the mixture was spotted onto Potato Dextrose (PD) agar plates supplemented with 1% activated charcoal. Plates were incubated for 18 h at 25°C. Mating was scored by the appearance of white, fuzzy aerial hyphae. All assays were conducted in triplicate (biological replicates), with at least three technical replicates per experiment.

For growth curve assays, cells were cultured in 96-well plates under constant shaking at 28ºC. The $OD_{600}$ was monitored every 15 min using a Spark 10M (Tecan) fluorescence microplate reader. Three biological replicates, each including three technical replicates, were performed until cultures reached an $OD_{600}$ of 1.0–1.2.

#### Flow Cytometry

To analyze the DNA content of the Δ*lem2* mutant, we employed a modified protocol (9). Cells were grown in YEPSL medium overnight at 28°C, diluted into fresh YEPSL medium, and cultured until reaching an $OD_{600}$ of 0.6–0.8. Before analysis, cells were harvested by centrifugation, washed twice with cold sterile water and fixed in 70% ethanol overnight at 4°C. Following fixation, cells were resuspended in 50 mM sodium citrate (pH 7.5). Cellular RNA was degraded with RNase A (0.25 mg/mL) for 1 h at 50°C, followed by proteinase K (1 mg/mL) for an additional 1h at 50°C to remove proteins.

Finally, DNA was stained with Propidium Iodide (PI) at a final concentration of 16 µg/mL. The fluorescence of 10,000 cells was measured using a FACSCalibur flow cytometer (Becton Dickinson) to determine the distribution of 1C and 2C DNA content.

**Plant Infection Assays**

To evaluate the virulence of the different *U. maydis* strains, pathogenicity assays were conducted using *Zea mays* (cv. Early Golden Bantam). Fungal cultures were grown in YEPSL medium to exponential phase ($OD_{600}$ 0.6–0.8). Cells were harvested by centrifugation, washed twice with sterile distilled water, and resuspended to a final density of $OD_{600}$ 3.0 (except for chlorazol black staining, where an $OD_{600}$ of 1.0 was used). Seedlings were inoculated seven days after sowing by injecting the cell suspension into the base of the stem. Disease progression and tumor development were scored 14 days post-infection (dpi). Each assay was performed in three independent biological rounds. To determine statistical significance, the ordinal logistic mixed model (GLMM) was applied, with a significance threshold of $p < 0.05$.

**Sample preparation for microscopy**

For immobilization during live-cell imaging, cells were placed on 2% agarose pads prepared with the appropriate medium: YEPSL for vegetative growth, MM-NO3 for filamentous growth, or CMU for conjugation tube assays. To visualize filamentous growth in AB33 strains, cultures were grown in YEPSL to an $OD_{600}$ of 0.5–0.7. Filamentation was induced by shifting the cultures to MM-NO3 (supplemented with 1% glucose) and incubating for 3 h at 28°C, unless otherwise indicated. Subsequently, cells were transferred onto the agarose pads and further incubated for 30–60 min at 28°C to allow for stabilization prior to image acquisition.

Live-cell imaging was performed using various microscopy systems tailored to specific experimental requirements. For general localization assays, optical sections were acquired at 0.4 µm intervals using a Spinning Disk confocal microscope. The co-expression of Lem2::GFP and NLS::3xRFP was analyzed using a DeltaVision microscope system, which allowed for the simultaneous acquisition of fluorescence and differential interference contrast (DIC) images. To assess nuclear DNA content, filaments induced for 5 h were stained with DAPI.

To analyze *U. maydis* progression within maize tissue, infected leaves were harvested at 3 and 5 days post-inoculation (dpi) and processed as previously described (10). Samples were cleared in ethanol, treated with 10% KOH at 60°C for 4 h, and washed in 1x PBS. Fungal hyphae were stained with WGA-Alexa Fluor 488 (green), while plant cell walls were stained with Propidium Iodide (red). Additionally, fungal structures at 1 dpi were visualized using chlorazole black staining as previously described (11). Images were acquired using the DeltaVision microscope system. All microscopy images were processed and analyzed using ImageJ/Fiji software.

To assess telomere distribution, the nuclear area in each optical section was divided into three concentric zones of equal area, as previously described (12). These regions included the nuclear periphery (Zone I), the intermediate region (Zone II), and the nuclear interior (Zone III). The distance between telomeric foci and the nuclear envelope was determined in ImageJ/Fiji by performing a line profile analysis. A line was drawn across the nucleus to generate a fluorescence intensity profile, which allowed for the precise identification of peaks corresponding to the telomere and the nuclear envelope signals. The measured distance between these peaks was then used to assign each telomere to its respective zone.

**Microscopy Systems and Settings**

For high-resolution imaging, the DeltaVision System (Applied Precision) was utilized, based on an Olympus IX71 inverted microscope equipped with a CoolSnap HQ camera and an environmental control chamber for temperature and $CO_2$ regulation. This system supported both brightfield and Differential Interference Contrast (DIC) imaging. Fluorescence was detected using specific filter sets, including GFP (Excitation 470/40, Emission 528/38), DsRed/mCherry (Excitation 555/28, Emission 617/73), and DAPI (Excitation 360/40, Emission 457/50). Additionally, a Spinning Disk Confocal system (Roper Scientific) was employed, featuring an Olympus IX81 inverted microscope coupled to a Yokogawa CSU-X1 spinning disk unit. Image acquisition was captured via CoolSnap HQ2 or Evolve cameras, with precise temperature control maintained by a Bioptechs FCS2 Perfusion System. Data acquisition and processing were managed through MetaMorph software. Simultaneous detection of GFP and mCherry was achieved using a Yokogawa CSU-W1 unit equipped with 488 nm and 561 nm lasers.

#### Immunofluorescence of Tubulin and Actin

For immunofluorescence analysis, *U. maydis* cells were grown in YEPSL to an $OD_{600}$ of 0.6–0.8, diluted to 0.05, and harvested at a final $OD_{600}$ of 0.5–0.6. Cells were processed as previously described (13), but omitting the methanol treatment. Cell wall degradation was achieved using the Lalzyme enzyme complex. Tubulin was detected using monoclonal α-tubulin antibody (Sigma-Aldrich, produced in mouse), while actin was identified with monoclonal anti-actin mouse antibody (clone AC-40). Both primary antibodies were subsequently visualized by incubation with a common Alexa Fluor 488-conjugated anti-mouse antibody. Images were acquired using the Spinning Disk confocal microscope and processed with ImageJ/Fiji.

#### RNA-seq analysis

Total RNA was submitted to Tech Solutions (Hong Kong) Co., Limited. Library preparation and paired-end sequencing were performed by BGI using the DNBseq PE100 RNA-seq platform. Trimmed reads provided by BGI were mapped to the *U. maydis* genome using HISAT2 (v2.2.1) (95) with strand-specific settings (reverse). The resulting SAM files were converted to sorted and indexed BAM files using Samtools (v1.17) (14). Gene-level counts were generated with HTSeq-count (v2.0.5) (96), counting reads overlapping annotated gene features (GFF3 annotation SOURCE). Prior to downstream analysis, genes with low counts (<10 reads in at least half of the samples in any group) were filtered. Differential expression analysis was performed using DESeq2 (v1.38) (97), incorporating replicate effects into the design. Genes with an adjusted $P_{adj}$ <0.05 and $log_2FC \geq 0.5$ were considered significantly differentially expressed. For these genes, variance-stabilizing transformed (VST) counts derived from the DESeq2 model were averaged by condition, scaled by gene (z score), and clustered using k-means to identify expression patterns. Gene Ontology (GO) biological process and KEGG pathway (15) enrichment analyses were performed using ShinyGO 0.80 (16), with a significance threshold of FDR < 0.05.

**Table S1** Strains used in this study

| Strain | Genotype | Resistance | Source | Strain ID |
|---|---|---|---|---|
| SG200 | *a1 mfa2 bE1 bW2* | - | (17) | 17 |
| SG200 Δ*lem2* | *a1 mfa2 bE1 bW2* Δ*lem2* | *NatR* | This work | 416 |
| SG200 Δ*lem2-lem2* | *a1 mfa2 bE1 bW2* Δ*lem2 ipR*[*Potef:lem2*]*ipS* | *NatR CbxR* | This work | 1581 |
| SG200 *lem2*::GFP | *a1 mfa2 bE1 bW2 lem2*::GFP | *HygR* | This work | 944 |
| SG200 *lem2*::GFP NLS::3xRFP | *a1 mfa2 bE1 bW2 lem2*::GFP *ipR*[*Potef*:NLS::3xRFP]*ipS* | *HygR CbxR* | This work | 1201 |
| SG200 NLS::3xRFP *nup214*::GFP | *a1 mfa2 bE1 bW2 ipR*[*Potef*:NLS::3xRFP]*ipS nup214*::GFP | *CbxR HygR* | This work | 963 |
| SG200 Δ*lem2* NLS::3xRFP *nup214*::GFP | *a1 mfa2 bE1 bW2* Δ*lem2 ipR*[*Potef*:NLS::3xRFP]*ipS nup214*::GFP | *NatR CbxR HygR* | This work | 983 |
| SG200 *nup214*::mCherry *rap1*::GFP | *a1 mfa2 bE1 bW2 rap1*::GFP *nup214*::mCherry | *NatR HygR GenR* | This work | 1527 |
| SG200 Δ*lem2* *nup214*::mCherry *rap1*::GFP | *a1 mfa2 bE1 bW2* Δ*lem2 rap1*::GFP *nup214*::mCherry | *NatR HygR GenR* | This work | 1525 |
| AB33 | *a2* P*nar:bW2 bE1* | - | (18) | 855 |
| AB33 Δ*lem2* | *a2* P*nar:bW2 bE1* Δ*lem2* | *NatR* | This work | 909 |
| AB33 *nup214*::mCherry *mis12*::GFP | *a2* P*nar:bW2 bE1 mis12*::GFP *nup214::*mCherry | *HygR GenR* | This work | 1380 |
| AB33 Δ*lem2* *nup214::mCherry* *mis12*::GFP | *a2 Pnar:bW2 bE1* Δ*lem2 mis12*::GFP *nup214*::mCherry | *NatR HygR GenR* | This work | 1386 |
| AB33 *nup214*::mCherry *grc1*::GFP | *a2* P*nar:bW2 bE1 grc1*::GFP *nup214::*mCherry | *HygR GenR* | This work | 1376 |
| AB33 Δ*lem2* *nup214*::mCherry *grc1*::GFP | *a2* P*nar:bW2 bE1* Δ*lem2 grc1*::GFP *nup214*::mCherry | *NatR HygR GenR* | This work | 1382 |
| AB33 *nup214*::mCherry *rap1*::GFP | *a2* P*nar*:*bW2 bE1 rap1*::GFP *nup214*::mCherry | *HygR GenR* | This work | 1498 |
| *AB33* Δ*lem2* *nup214::mCherry* *rap1*::GFP | *a2* P*nar: bW2 bE1* Δ*lem2 rap1*::GFP *nup214*::mCherry | *NatR HygR GenR* | This work | 1500 |
| FB1 | *a1 b1* | - | (19) | 14 |
| FB2 | *a2 b2* | - | (19) | 15 |
| FB1 Δ*lem2* | *a1 b1* Δ*lem2* | *NatR* | This work | 430 |
| FB2 Δ*lem2* | *a2 b2* Δ*lem2* | *NatR* | This work | 435 |
| SG200 AM1::GFP | *a2 b2* P01779:*am1*::GFP | *GenR* | (20) | 187 |
| SG200 Δ*lem2* AM1::GFP | *a2 b2* Δ*lem2* P01779:*am1*::GFP | *NatR GenR* | This work | 884 |
| AB33 *nup214*::GFP NLS::3xRFP | *a2* P*nar*:*bW2 ipR*[*Potef*:NLS::3xRFP]*ipS nup214*::GFP | *CbxR HygR* | This work | 1021 |
| AB33 Δ*lem2* *nup214*::GFP *NLS*::3xRFP | *a2* P*nar:bW2* Δ*lem2 ipR*[*Potef*:NLS::3xRFP]*ipS nup214*::GFP | *NatR CbxR HygR* | This work | 1060 |

| | | | | |
|---|---|---|---|---|
| AB33 *nup214*::mCherry *pot1*::GFP | *a2* Pnar: *bW2 pot1*::GFP *nup214*::mCherry | *HygR GenR* | This work | 1378 |
| AB33 Δ*lem2-lem2* *nup214*::mCherry *rap1*::GFP | *a2* Pnar: *bW2* Δ*lem2* *ipR*[Potef:*lem2*]*ipS* *nup214*::mCherry *rap1*::GFP | *NatR Cbx GenR HygR* | This work | 1556 |
| AB33 Δ*lem2-LEM* *nup214*::mCherry *rap1*::GFP | *a2* Pnar*: bW2* Δ*lem2* *ipR*[Potef*:LEM*]*ipS* *nup214::*mCherry *rap1*::GFP | *NatR Cbx GenR HygR* | This work | 1558 |
| AB33 Δ*lem2-MSC* *nup214*::mCherry *rap1*::GFP | *a2* Pnar*: bW2* Δ*lem2* *ipR*[Potef*:MSC*]*ipS* *nup214*::mCherry *rap1*::GFP | *NatR Cbx GenR HygR* | This work | 1559 |
| SG200 Δ*lem2-LEM* | *a1 mfa2 bE1 bW2* Δ*lem2* *ipR*[Potef*:LEM*]*ipS* | *NatR CbxR* | This work | 1582 |
| SG200 Δ*lem2-MSC* | *a1 mfa2 bE1 bW2* Δ*lem2* *ipR*[Potef*:MSC*]*ipS* | *NatR CbxR* | This work | 1584 |
| AB33 *nup214*::mCherry *rap1*::GFP *nup84*::GBP | *a2* Pnar:*bW2 bE1 rap1*::GFP *nup214*::mCherry *nup84*::GBP | *HygR GenR CbxR* | This work | 1513 |
| AB33 Δ*lem2* *nup214*::mCherry *rap1*::GFP *nup84*::GBP | *a2* Pnar:*bW2 bE1* Δ*lem2* *rap1*::GFP *nup214*::mCherry *nup84*::GBP | *NatR HygR GenR CbxR* | This work | 1512 |
| SG200 *nup214*::mCherry *rap1*::GFP *nup84*::GBP | *a1 mfa2 bE1 bW2 rap1*::GFP *nup214*::mCherry *nup84*::GBP | *HygR GenR CbxR* | This work | 1541 |
| SG200 Δ*lem2* *nup214*::mCherry *rap1*::GFP *nup84*::GBP | *a1 mfa2 bE1 bW2* Δ*lem2* *rap1*::GFP *nup214*::mCherry *nup84*::GBP | *NatR HygR GenR CbxR* | This work | 1543 |

**Table S2** Plasmids used in this study

| Plasmid Cloning | Cloning method | Primer Name | Primer sequence 5'-3' |
|---|---|---|---|
| pGEMT Δ*um00208-natR* | Standard molecular cloning | COum00208-5 | AGCATCTTCCAATCCTCGTTCC |
| | | COum00208-3 | CCAAAGAAGAACTGAGGTAGGAGG |
| | | um00208KO5-1 | CTCATCTACTCGCTTCGGAGG |
| | | um00208KO5-2 | CACGGCCTGAGTGGCCGCTTCATACAAACACACCTGTCG |
| | | um00208KO3-1 | GTGGCCATCTAGGCCCAAGGAAATCACCAGGACAACG |
| | | um00208KO3-2 | ATGGCTAGAGAGAAAGGATACG |
| p123 P*otef*::*lem2* | Standard molecular cloning | Nco1_Lem2_F | ATACCATGGGTCGAACGCGACGAG |
| | | Not1_Lem2_R | ATAGCGGCCGCTTAAGCAACAGGCATCC |

| | | | |
|---|---|---|---|
| p123 P*otef*::*lem2_MSC+TM* | Standard molecular cloning | Nco1_Lem2-MSC-F | ATACCATGGCCAAGTCTTT |
| | | Not1_Lem2_R | ATAGCGGCCGCTTAAGCAACAGGCATCC |
| p123 P*otef*::lem2_*LEM+TM* | Standard molecular cloning | Nco1_Lem2_F | ATACCATGGGTCGAACGCGACGAG |
| | | Not1-Lem2-Heh-R | ATAGCGGCCGCTTATCGCAAGTAGAAGAATCC |
| pRS426 *Um00208*::GFP-*HygR* | Standard molecular cloning | GFP-h-MESC-5' | ATGGTGAGCAAGGGCGAGGA |
| | | GFP-h-MESC-3' | ATAGGGCGAATTGGAGCTCG |
| | | Lem2_UP_MESC-1 | TCGAGGTCGACGGTATCGATAAGCTTGATACGCATACGGAAAAGTACTCG |
| | | Lem2_DOWN_MESC-2 | GCTCTAGAACTAGTGGATCCCCCGGGCTGCACAGCTCAATCTCTCTATGC |
| | | Lem2_UP_MESC-2 | GGTGAACAGCTCCTCGCCCTTGCTCACCATAGCAACAGGCATCCTCGCAT |
| | | Lem2_DOWN_MESC-1 | CCTGAGTGGCCGAGCTCCAATTCGCCCTATTGGTTGCACACCAGAAGCAC |
| pC_NLS::3xRFP-*CbxR* (P*otef*-gal4s-mrfp-mrfp-mrfp) | (7) | | |
| pN214G (*nup214*-GFP, *HygR*) | (8) | Fwd UM_01089 | GCTCTAGCAATACGGTCAATG |
| | | Rev UM_01089 | GCAAATTTGATGCCGAGATCT |
| pBSK *nup214*::mCherry-*GenR* | NEBuilder® HiFi DNA Assembly | Nup214-5_fwd (tagging Nucleoporin red) | cgaattcctgcagcccggggTGCAGCCGTTGACGACAAG |
| | | Nup214-5_rev (tagging Nucleoporin red) | tgctcaccatGTCCGATTTGTCCACGTTCC |
| | | mCherry-gen_fwd (tagging Nucleoporin red) | caaatcggacATGGTGAGCAAGGGCGAG |
| | | mCherry-gen_rev (tagging Nucleoporin red) | gatgatgagaGCAAATTAAAGCCTTCGAGCG |
| | | Nup214-3_fwd (tagging Nucleoporin red) | tttaatttgcTCTCATCATCCTTTCTCCAATCG |
| | | Nup214-3_rev (tagging Nucleoporin red) | cggccgctctagaactagtgAAGCGACCCACGGTCATC |
| | | Ampl Nup214cherry Fwd | TGCAGCCGTTGACGACAAG |
| | | Ampl Nup214cherry rev | AAGCGACCCACGGTCATC |
| | | Fw_compmCherry-Nup214 | GACCCTTCCACATTCAGTT |
| | | Rev_compmCherry-Nup214 | CTCTCATTCAAGACATCGT |
| pBSK *rap1*::GFP-*HygR* | NEBuilder® HiFi DNA Assembly | TagRap1KO5NEB_Fw | cgaattcctgcagcccggggTTTCTACGACTTCACCATGG |
| | | TagRap1KO5NEB_rev | tgctcaccatTCCACGGACTCGCTTTAAG |

| | | | |
|---|---|---|---|
| | | TagRap1_GFPHyg_fwNEB | agtccgtggaATGGTGAGCAAGGGCGAG |
| | | TagRap1__GFPHygNEB_rev | aatcaagcgcTATTAATGCGGCCGCACAG |
| | | TagRap1_KO3NEB_Fw | cgcattaataGCGCTTGATTTGCTCCTC |
| | | TagRap1_KO3NEB_rev | cggccgctctagaactagtgTATCGACCTTCCAGCGTTTC |
| | | Rap1ko5_sacarinserto_Fw | TTTCTACGACTTCACCATGG |
| | | Rap1ko3_sacarinserto_Rev | TATCGACCTTCCAGCGTTT |
| | | sec_Ust_Rap__KO5_Fw | GCTCATCAGCAAAGTTACTG |
| | | sec_Ust_Rap__KO5_rev | CCACGGACTCGCTTTAAGG |
| | | sec_Ust_Rap__KO3_Fw | TGTCTGTTAAAGTCACCTTG |
| | | sec_Ust_Rap__KO3_rev | ATCATTTGCACTTCGGCAA |
| pBSK *mis12*::3xGFP-*HygR* | NEBuilder® HiFi DNA Assembly | Mis12-5_fwd (tagging kinetochore) | CGAATTCCTGCAGCCCGGGGCATCGCTACTGAGTTCGAGAATGTAGTG |
| | | Mis12-5_rev (tagging kinetochore) | TGCTCACCATGCCTCGAGAGCGAGTCGC |
| | | Fwd_GFP-HygR_(Mis12)_NEB_new | CTCTCGAGGCAACGCGGCCACCATGGTGA |
| | | 3xGFP::HygR_rev (tagging kinetochore) | AATAAAGGGTTGCGGCCGCACAGCTTCG |
| | | Mis12-3_fwd (tagging kinetochore) | TGCGGCCGCAACCCTTTATTCTGTATTAAAACTTG |
| | | Mis12-3_rev (tagging kinetochore) | CGGCCGCTCTAGAACTAGTGCCAGGAGATTCTCGAGATC |
| | | Fwd_GFP-HygR_(Mis12)_NEB_new | CTCTCGAGGCAACGCGGCCACCATGGTGA |
| | | Ampl Mis12GFP Fwd | CATCGCTACTGAGTTCGAGAATGTAGTG |
| | | Ampl Mis12GFP Rev | CCAGGAGATTCTCGAGATC |
| pBSK *grc1*::3xGFP-*HygR* | NEBuilder® HiFi DNA Assembly | Grc1-5_fwd (tagging SPB) | CGAATTCCTGCAGCCCGGGGACAGTCGGCACATGAGCTG |
| | | Grc1-5_rev (tagging SPB) | TGCTCACCATCAAATTCGAGCTCTGAAACCTG |
| | | Fwd_GFPHygR_(Grc1)_NEB_new | CTCGAATTTGAACGCGGCCACCATGGTGA |
| | | 3xGFP-Hyg_rev (tagging SPB) | GACTCGACAGTGCGGCCGCACAGCTTCG |
| | | Grc1-3_fwd (tagging SPB) | TGCGGCCGCACTGTCGAGTCATCCAGCC |
| | | Grc1-3_rev (tagging SPB) | CGGCCGCTCTAGAACTAGTGCATGCTGCTTGACGCAATC |
| | | Grc1-GFP_Ampli FWD | ACAGTCGGCACATGAGCTGG |

| | | | |
|---|---|---|---|
| | | Grc1-GFP_Ampli Rev | CATGCTGCTTGACGCAATCTCC |
| p*Pot1*::3xGFP-*HygR* | NEBuilder® HiFi DNA Assembly | Pot1-5_fwd (tagging telomeres) | CGAATTCCTGCAGCCCGGGGACAAGCGCAGCAGGGTGC |
| | | Pot1-5_rev (tagging telomeres) | TGCTCACCATCAATAGATCGTGTTCGTCAGATAGAACGTTG |
| | | Fwd_GFPHygR_(Pot1)_NEB_new | CGATCTATTGAACGCGGCCACCATGGTGA |
| | | 3xGFP-HygR_rev (tagging telomeres) | CCCACCGACTTGCGGCCGCACAGCTTCG |
| | | Pot1-3_fwd (tagging telomeres) | TGCGGCCGCAAGTCGGTGGGCCAGACCC |
| | | Pot1-3_rev (tagging telomeres) | CGGCCGCTCTAGAACTAGTGGCTGCGGCAGGTGCGATTG |
| | | Pot1-GFP_Ampli FWD | ACAAGCGCAGCAGGGTGC |
| | | Pot1-GFP_Ampli Rev | GCTGCGGCAGGTGCGATTG |
| pBSK *nup84*::GBP::*nos*:*CbxR* | NEBuilder® | KO5_Nup84_fwd | cgaattcctgcagcccgggggtcatgctacgtacaggtattg |
| | | KO5_Nup84_rev | ccagctgcaccgattctacagccttctg |
| | | ko3_Nup84_fwd | gatccccattctgcaccatcgcatctgataac |
| | | ko3_Nup84_rev | cggccgctctagaactagtgggcttcgatgatttagtgctg |
| | | Nup84_GFPnanobody_fwd | tgtagaatcggtgcagctggtggagtctg |
| | | Nup84_Cbx-Kassete_rev | gatggtgcagaatggggatcttcgctcaac |
| | | Fwd sacar inserto Nup84 | GTCATGCTACGTACAGGTA |
| | | Rev Sacar insert Nup84 | GGCTTCGATGATTTAGTGC |
| | | Nup84 comprobar Fwd | CATCTGATCCTCTACCTCC |
| | | Nup84 comprobar Rev | GCTGAAATGTCTCCTTCAAC |

**Table S3** Gene numbers

| Protein ID | Gene number (UMAG) |
|---|---|
| Lem2 | UMAG_00208 |
| Rap1 | UMAG_04676 |
| Nup84 | UMAG_04795 |
| GRC1 | UMAG_11816 |
| Mis12 | UMAG_04180 |
| Pot1 | UMAG_05117 |
| Nup214 | UMAG_01089 |

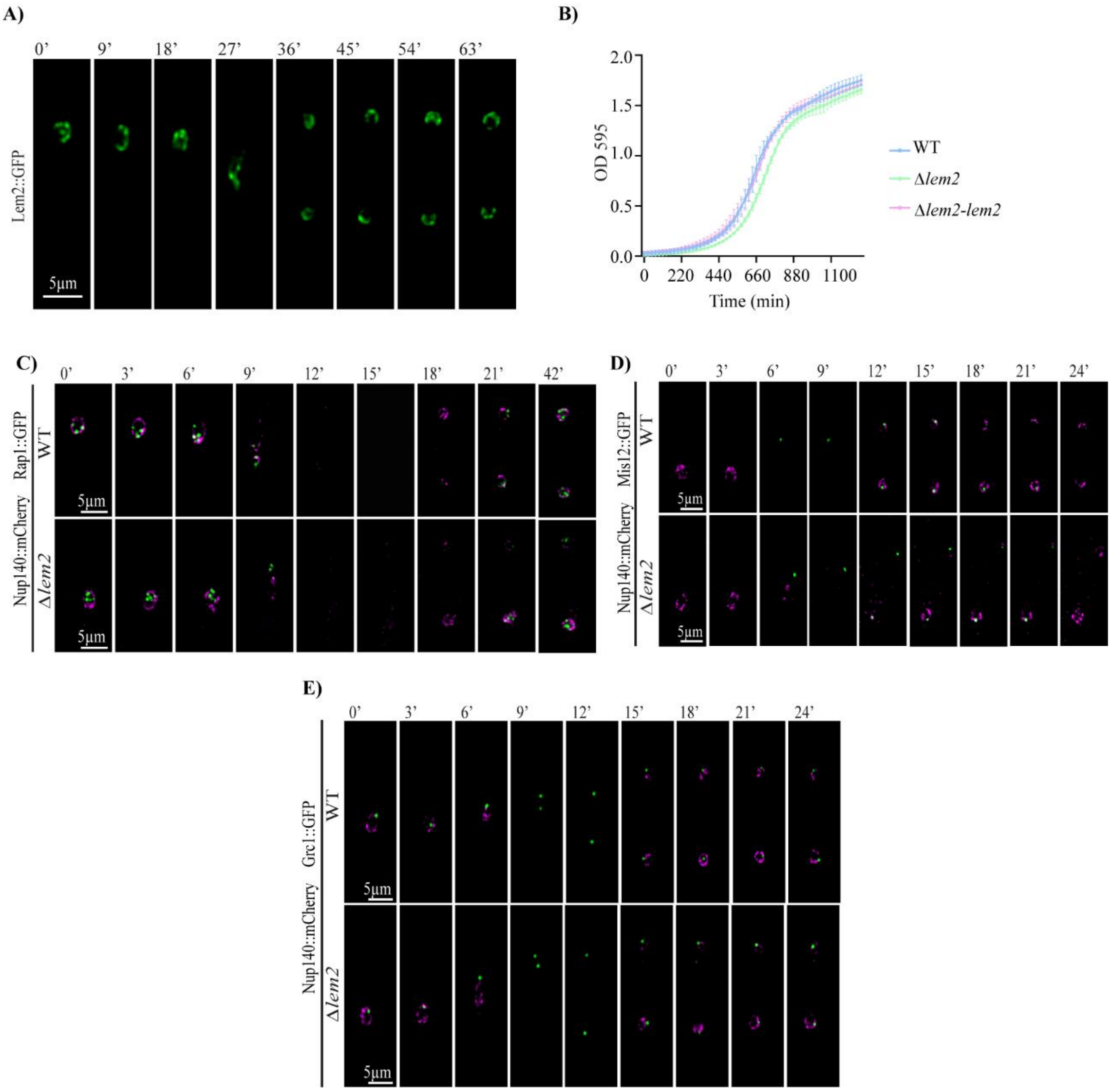


**Figure S1. Mitotic localization of Lem2, telomeres, and centromeres**. **(A)** Time-lapse microscopy of a cell showing the dynamic distribution of Lem2::GFP. **(B)** Growth curves of WT, Δ*lem2*, and the Δ*lem2-lem2* strains. Error bars represent the standard deviation (SD) from three independent replicates. **(C–E)** Representative time-lapse images of WT and Δ*lem2* cells expressing Nup214::mCherry (magenta) and the following GFP-tagged proteins (green): **(C)** the telomere-binding protein Rap1::GFP, **(D)** the inner kinetochore protein Mis12::GFP, and **(E)** the centromere-associated protein Grc1::GFP. Numbers indicate time in minutes. Images are maximum intensity projections of Z-stacks. Panels A and B use the SG200 background; panels C and D, the AB33 background.

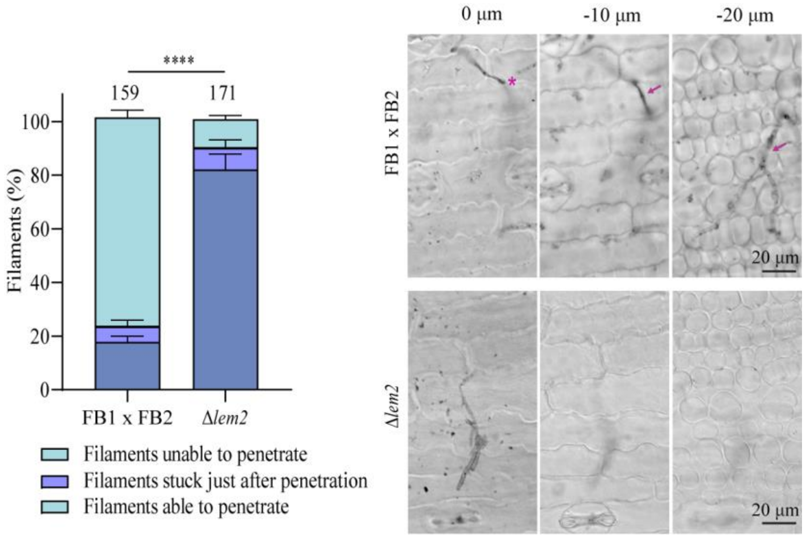


**Figure S2. Analysis of appressorium penetration.** Maize leaves infected with FB1×FB2 and FB1 Δ*lem2* × FB2 Δ*lem2* were stained with chlorazol black at 24 hours post-inoculation (hpi). Left panel: Maximum intensity projections showing appressorium formation (magenta asterisks) and invasive hyphae (magenta arrows). Right panel: Quantification of infection stages for each strain. Error bars represent the SD from three independent replicates. Statistical significance was determined using the ordinal logistic mixed model (GLMM) (****, P < 0.0001). The total number of filaments analyzed is indicated above each column.

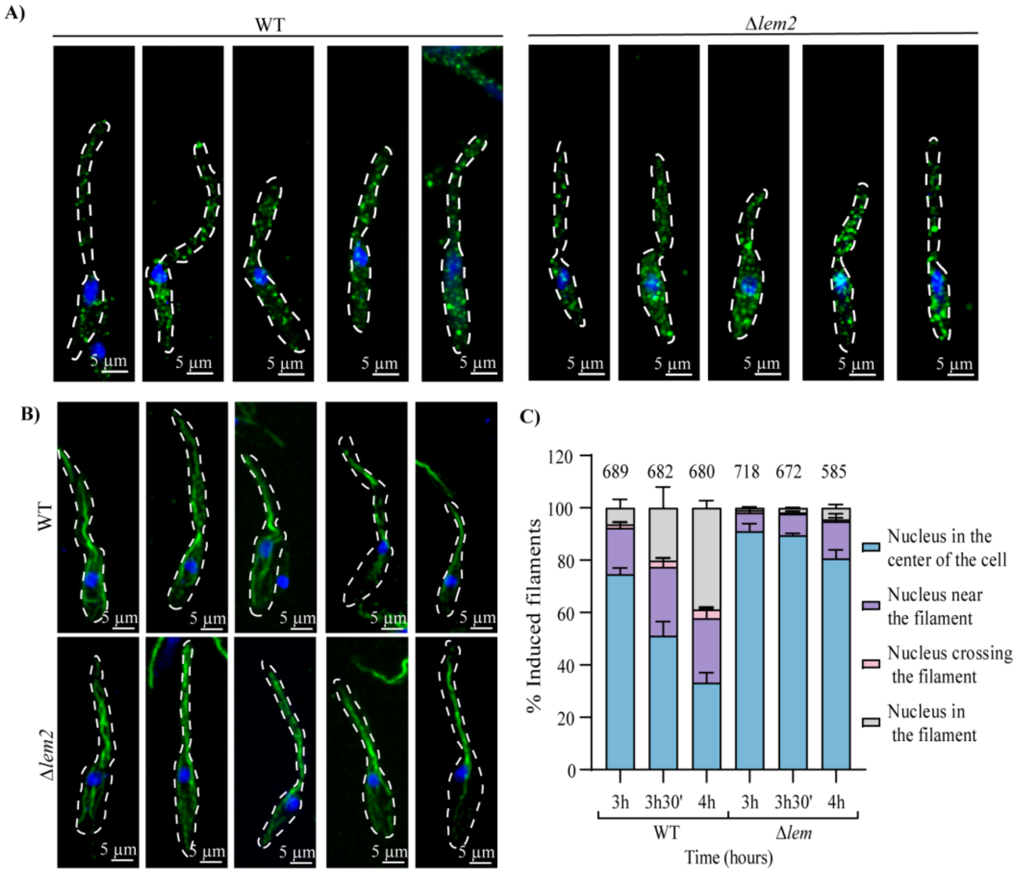


**Figure S3. Cytoskeleton organization and nuclear migration kinetics**. **(A–B)** Immunofluorescence staining of actin (green) **(A)** and α-tubulin (green) **(B)** in AB33 WT and Δ*lem2* filaments at 3.5 hpi. Nuclei were counterstained with DAPI (blue). **(C)** Quantification of nuclear position at the indicated time points after induction of filamentation in AB33 WT and Δ*lem2* strains. Error bars represent the SD from three independent replicates. The total number of filaments analyzed (n) is indicated above each column.

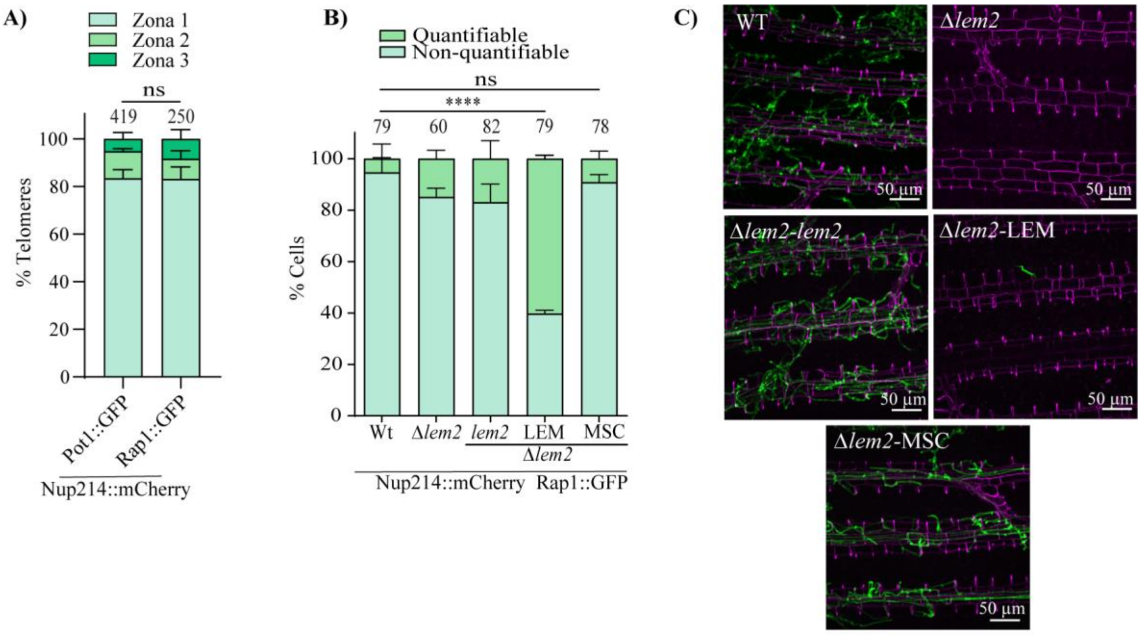


**Figure S4. Telomere distribution patterns and pathogenicity assays**. **(A)** Quantification of telomere distribution relative to the nuclear periphery (Zones I–III) in WT cells expressing Nup214:mCherry and telomere markers, either Rap1::GFP or Pot1::GFP. **(B)** Proportion of cells with quantifiable vs. non-quantifiable telomere localization in the indicated strains. Non-quantifiable cells represent those with compromised nuclear envelope integrity. **(C)** Representative images of maize leaves infected with the indicated strains, stained with WGA-AF488 (green) and Propidium Iodide (magenta) to visualize fungal colonization. In panels (A, B), error bars represent the SD from three independent replicates. Statistical significance was determined using the ordinal logistic mixed model (GLMM) (ns, not significant; ****, $P < 0.0001$). The total number of cells or filaments analyzed (*n*) is indicated above each column. Panels A and B show cells in the AB33 genetic background, while panel C uses the SG200 background.

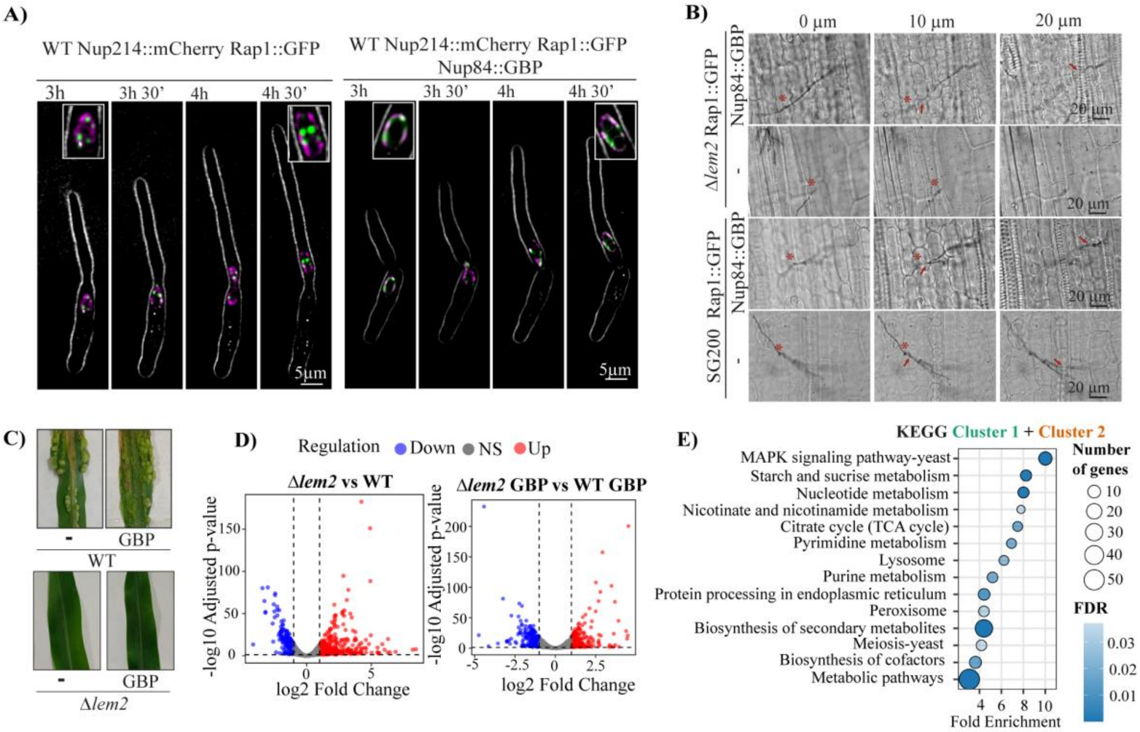


**Figure S5. Artificial telomere anchoring during pathogenesis and transcriptomic analysis.** **(A)** Representative images of nuclear positioning during filamentation in the indicated strains. Numbers above each frame indicate the time in minutes after induction of the filamentation. **(B)** Maize leaves infected with indicated strains and stained with chlorazol black at 24 hpi. Maximum intensity projections show representative filaments for each strain; red asterisks indicate appressorium formation and red arrows highlight invasive hyphae. The total number of plants analyzed is indicated above each column. **(C)** Maize leaves infected with the indicated strains at 14 dpi showing disease symptoms and tumor formation. **(D)** Volcano plots of differentially expressed genes (DEGs). Transcriptomic profiles at 3.5 hpi between are compared between: (left) Δ*lem2* (Nup214::mCherry Rap1::GFP) relative to WT (Nup214::mCherry Rap1::GFP); and (right) Δ*lem2* artificial anchoring strain (Nup214::mCherry Rap1::GFP Nup84::GBP) relative to its respective WT (Nup214:mCherry Rap1:eGFP Nup84:GBP). Red and blue dots represent significantly upregulated ($\log_2$FC > 1) and downregulated ($\log_2$ fold change < -1) genes, with $P_{adj}$ < 0.05. **(E)** Functional enrichment of KEGG pathways significantly enriched among genes from Cluster 1 and Cluster 2. Analysis was performed using ShinyGO 0.85 (FDR < 0.05). All strains used in this figure express Nup214::mCherry and Rap1::GFP. Panels A, D, and E show cells in the AB33 genetic background, while panels B and C use the SG200 background.

### References


1. Sambrook, J., Fritsch, E. R., & Maniatis, T. (1989). Molecular Cloning A Laboratory Manual (2nd ed.). Cold Spring Harbor, NY Cold Spring Harbor Laboratory Press. - References - Scientific Research Publishing.

2. C. Aichinger, *et al.*, Identification of plant-regulated genes in Ustilago maydis by enhancer-trapping mutagenesis. *Molecular Genetics and Genomics 2003 270:4* **270**, 303–314 (2003).

3. B. Gillissen, *et al.*, A two-component regulatory system for self/non-self recognition in Ustilago maydis. *Cell* **68**, 647–657 (1992).

4. A. Brachmann, J. König, C. Julius, M. Feldbrügge, A reverse genetic approach for generating gene replacement mutants in Ustilago maydis. *Molecular genetics and genomics : MGG* **272**, 216–226 (2004).

5. K. Bösch, *et al.*, Genetic Manipulation of the Plant Pathogen Ustilago maydis to Study Fungal Biology and Plant Microbe Interactions. *Journal of visualized experiments : JoVE* **2016** (2016).

6. J. P. R. Keon, G. A. White, J. A. Hargreaves, Isolation, characterization and sequence of a gene conferring resistance to the systemic fungicide carboxin from the maize smut pathogen, Ustilago maydis. *Current genetics* **19**, 475–481 (1991).

7. A. Straube, I. Weber, G. Steinberg, A novel mechanism of nuclear envelope break-down in a fungus: nuclear migration strips off the envelope. *The EMBO journal* **24**, 1674–1685 (2005).

8. U. Theisen, A. Straube, G. Steinberg, Dynamic Rearrangement of Nucleoporins during Fungal "Open" Mitosis. *Molecular Biology of the Cell* **19**, 1230 (2008).

9. T. García-Muse, G. Steinberg, J. Pérez-Martín, Pheromone-induced G2 arrest in the phytopathogenic fungus Ustilago maydis. *Eukaryotic Cell* **2**, 494–500 (2003).

10. A. Redkar, E. Jaeger, G. Doehlemann, Visualization of Growth and Morphology of Fungal Hyphae in planta Using WGA-AF488 and Propidium Iodide Co-staining. (2018). https://doi.org/10.21769/BioProtoc.2942.

11. A. Brachmann, J. Schirawski, P. Müller, R. Kahmann, An unusual MAP kinase is required for efficient penetration of the plant surface by Ustilago maydis. *The EMBO Journal 2003 22:9* **22**, 2199–2210 (2003).

12. F. Hediger, A. Taddei, F. R. Neumann, S. M. Gasser, Methods for Visualizing Chromatin Dynamics in Living Yeast. *Methods in Enzymology* **375**, 345–365 (2004).

13. F. Banuett, A Method to Visualize the Actin and Microtubule Cytoskeleton by Indirect Immunofluorescence. *Methods in molecular biology (Clifton, N.J.)* **638**, 225–233 (2010).

14. H. Li, A statistical framework for SNP calling, mutation discovery, association mapping and population genetical parameter estimation from sequencing data. *Bioinformatics (Oxford, England)* **27**, 2987–2993 (2011).

15. M. Kanehisa, M. Furumichi, Y. Sato, M. Ishiguro-Watanabe, M. Tanabe, KEGG: integrating viruses and cellular organisms. *Nucleic Acids Res* **49**, D545–D551 (2021).

16. S. X. Ge, D. Jung, R. Yao, ShinyGO: a graphical gene-set enrichment tool for animals and plants. *Bioinformatics* **36**, 2628–2629 (2020).

17. M. Bölker, S. Genin, C. Lehmler, R. Kahmann, Genetic regulation of mating and dimorphism in Ustilago maydis. *Canadian Journal of Botany* **73**, 320–325 (1995).

18. A. Brachmann, G. Weinzierl, J. Kämper, R. Kahmann, Identification of genes in the bW/bE regulatory cascade in Ustilago maydis. *Molecular Microbiology* **42**, 1047–1063 (2001).

19. F. Banuett, I. Herskowitz, Different a alleles of Ustilago maydis are necessary for maintenance of filamentous growth but not for meiosis. *Proceedings of the National Academy of Sciences of the United States of America* **86**, 5878–5882 (1989).

20. A. Mendoza-Mendoza, *et al.*, Physical-chemical plant-derived signals induce differentiation in Ustilago maydis. *Molecular Microbiology* **71**, 895–911 (2009).
